# Evolutionary persistence of DNA methylation for millions of years after ancient loss of a *de novo* methyltransferase

**DOI:** 10.1101/149385

**Authors:** Sandra Catania, Phillip A. Dumesic, Harold Pimentel, Ammar Nasif, Caitlin I. Stoddard, Jordan E. Burke, Jolene K. Diedrich, Sophie Cook, Terrance Shea, Elizabeth Geinger, Robert Lintner, John R. Yates, Petra Hajkova, Geeta J. Narlikar, Christina A. Cuomo, Jonathan K. Pritchard, Hiten D. Madhani

**Author notes:** Contributed equally (Phillip A. Dumesic, Harold Pimentel).

## Abstract

Cytosine methylation of DNA is a widespread modification of DNA that plays numerous critical roles, yet has been lost many times in diverse eukaryotic lineages. In the yeast *Cryptococcus neoformans*, CG methylation occurs in transposon-rich repeats and requires the DNA methyltransferase, Dnmt5. We show that Dnmt5 displays exquisite maintenance-type specificity *in vitro* and *in vivo* and utilizes similar *in vivo* cofactors as the metazoan maintenance methylase Dnmt1. Remarkably, phylogenetic and functional analysis revealed that the ancestral species lost the gene for a *de novo* methylase, DnmtX, between 50-150 MYA. We examined how methylation has persisted since the ancient loss of DnmtX. Experimental and comparative studies reveal efficient replication of methylation patterns in *C. neoformans*, rare stochastic methylation loss and gain events, and the action of natural selection. We propose that an epigenome has been propagated for >50 MY through a process analogous to Darwinian evolution of the genome.

## INTRODUCTION

Methylation of cytosine on its fifth carbon (5mC) in DNA is found in all domains of life (Bird, 2002). The DNA of all known vertebrates and land plants harbor this modification, likely because of its critical function in genome defense (Zemach et al., 2010). 5mC is initially deposited by *de novo* DNA methyltransferases (DNMTs) that act on unmethylated substrates. In the context of palindromic sequence arrangements (e.g. CG and CHG) in which cytosines on both strands are methylated, this initial activity can then be “remembered” by maintenance enzymes that function on hemimethylated DNA produced by DNA replication (Law and Jacobsen, 2010). One example of this division of labor is found in the mammalian DNA methylation system. Here, *de novo* methylation is principally catalyzed by the Dnmt3a/b-Dnmt3L complex (Du et al., 2015). Once the DNA methylation mark is established during germline and embryonic development, it is then maintained by the Dnmt1 enzyme in association with UHRF1, a protein that recognizes H3K9me and hemimethylated 5mC (Bostick et al., 2007; Du et al., 2015; Law and Jacobsen, 2010). This division of labor between *de novo* and maintenance enzymes extends to land plants (Bronner et al., 2019), although Dnmt3 appears to have been lost in angiosperms and replaced by other *de novo* enzymes (Yaari et al., 2019). DNA methylation is essential for development and plays critical roles in silencing of transposable elements, chromosome stability, monoallelic gene expression, and gene silencing (Jones, 2012). Moreover, Dnmt3a is a commonly mutated driver gene in acute myeloid leukemia (Brunetti et al., 2017) and mutations in Dnmt3b cause the autosomal recessive disease ICF syndrome (Walton et al., 2014).

Despite having transposable elements, many animal, fungal, and protist species lack cytosine methylation, including the key model organisms *Saccharomyces cerevisiae*, *Schizosaccharomyces pombe*, *Caenorhabditis elegans*, *Drosophila melanogaster and Tetrahymena thermophila*. The corresponding ancestral species evidently lost the enzymes required for cytosine methylation. Supporting this conclusion, clades of insects and nematodes have been described that both contain orthologs of Dnmt3 and Dnmt1 and display genomic CG methylation (Bewick et al., 2017; Rosic et al., 2018). In ciliates, species harboring 5mC as well as those lacking it have been described (Wang et al., 2017). Likewise, the major divisions of the fungal kingdom, including ascomycetes and basidiomycetes, contain species with DNMTs and cytosine methylation, again suggesting that ancestral genomes contained 5mC (Bewick et al., 2019). Thus, 5mC appears to have been lost many times in various lineages. However, the underlying forces are unknown. More generally, analysis of genome sequences has led to the larger realization that gene loss is pervasive and a major force in evolution whose drivers are not well-understood (Albalat and Canestro, 2016).

The fungal kingdom offers rich phylogenetic diversity and many experimentally tractable systems. However, DNA methylation has only been extensively investigated in one species, in the filamentous ascomycete *Neurospora crassa* (Galagan and Selker, 2004). In this species, all stable 5mC is mediated by a single DNMT ortholog, DIM-2 (Aramayo and Selker, 2013). Based on the fact that cytosine methylation lost in DIM-2 knockout cells can be efficiently restored upon re-introduction of the gene (Galagan and Selker, 2004; Kouzminova and Selker, 2001), DIM-2 is thought to be a *de novo* enzyme, although enzymatic activity of purified DIM-2 has not been reported. DIM-2 and another cytosine methylase homolog, RID, promote repeat-induced point mutation (RIP) of duplicated sequences, a premeiotic process in which cytosine methylation and subsequent deamination produces C to T mutations (Gladyshev and Kleckner, 2017). The ability of *N. crassa* to convert transient cytosine methylation events to mutations may explain why this species seems to lack a maintenance methylase system. Classical studies of another ascomycete, *Ascobolus immersus*, provide evidence of memory mediated by DNA methylation.

In this species, duplicated sequences acquire cytosine methylation premeiotically, which is then propagated mitotically (Rossignol and Faugeron, 1995). This phenomenon predicts the existence of maintenance methylases in fungi. However, no such enzymes have been identified or biochemically characterized in this kingdom.

While nearly all metazoans and fungi that have DNA methylation encode at least two DNMT homologs in their genomes some, curiously, have only one (Bewick et al., 2019; Bewick et al., 2017; Rosic et al., 2018). One such species is the human fungal pathogen *Cryptococcus neoformans*, a basidiomycetous yeast that possesses symmetric CG methylation at transposon-rich centromeric and subtelomeric regions (Huff and Zilberman, 2014). This methylation is dependent on the predicted cytosine DNA methyltransferase (DNMT) Dnmt5, encoded by the *DMT5* gene (Huff and Zilberman, 2014). The Dnmt5 family is characterized by an N-terminal chromodomain (CD) followed by a cytosine methyltransferase catalytic domain, a RING finger, and a domain related to those of SNF2-type ATPases (Fig.1A) This putative enzyme is widespread in fungi and in green algae, several of which have been shown to have CG methylation that impacts nucleosome positioning (Huff and Zilberman, 2014). 5mC in *C. neoformans* is concentrated at transposable element-rich repeats; it likely functions in transposon silencing as a sister species, *C. deuterogattii*, has lost all active transposons and concomitantly lost 5mC and displays inactivating mutations in the *DMT5* gene (Yadav et al., 2018a).

**Figure 1.**
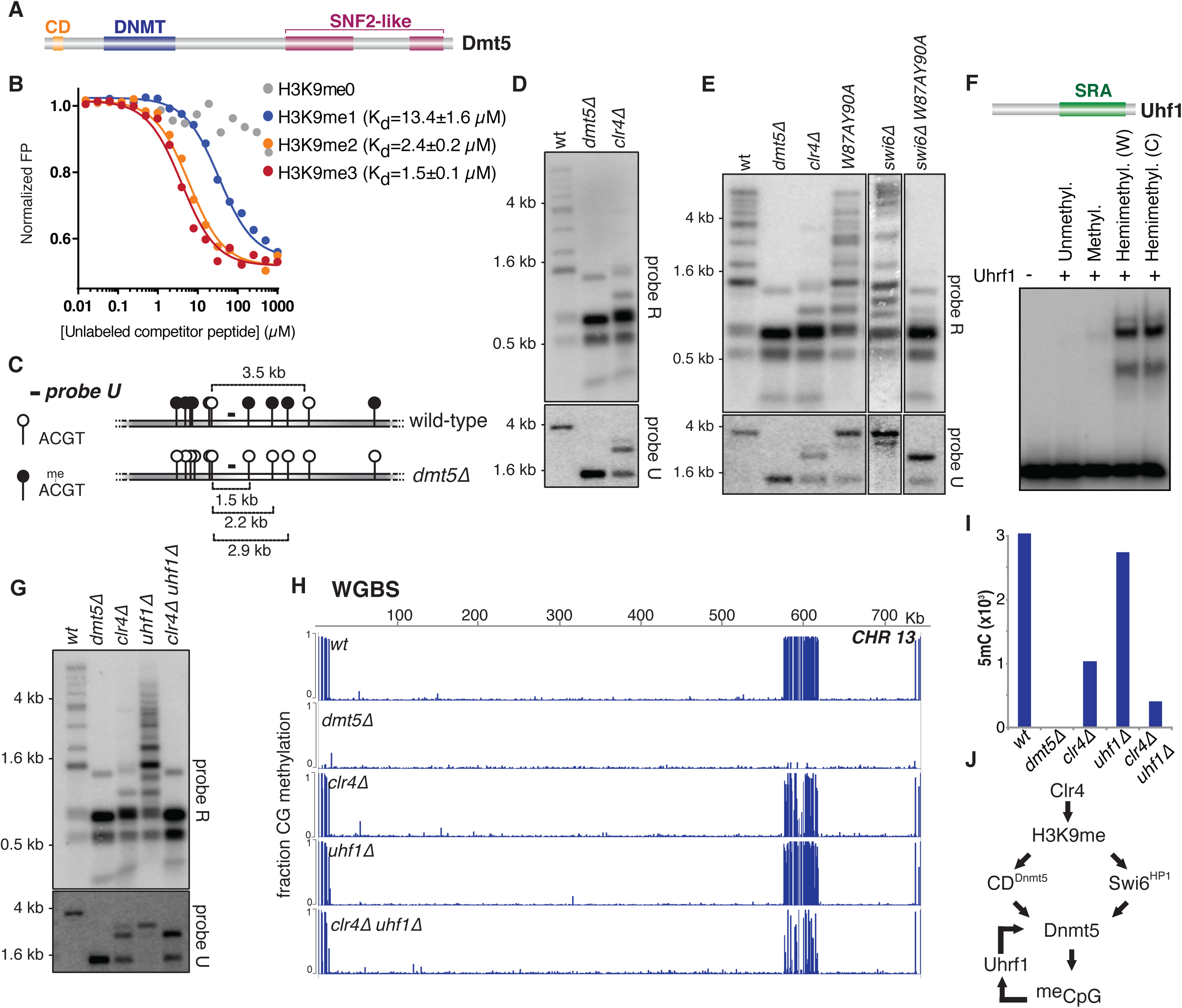
Multiple mechanisms promote efficient DNA methylation in *C. neoformans*. (A) Architecture of Dnmt5. CD: chromodomain; DNMT: DNA methyltransferase domain; SNF2-like: Swi/Snf ATPase domain. (B) Binding affinity of recombinant Dnmt5 CD to the indicated peptides as determined by competition fluorescence polarization. (C) Sites for the methylation-sensitive restriction endonuclease HpyCH4IV (ACGT) within a region of centromere 13. Filled circles: methylated sites; open circles: unmethylated sites. (D) 5mC levels of wild-type*, dmt5Δ* and *clr4Δ* assessed by Southern hybridization of HpyCH4IV-digested genomic DNA using probe corresponding to a repetitive sequence (probe R) or a unique sequence (probe U; panel B). (E) Southern analysis of 5mC in cells carrying CD mutant of Dnmt5 (*W87AY90A*), deletion of Swi6 (*swi6Δ)* or a combination of the two (*swi6*Δ *W87AY90A*). (F) Mobility shift assay testing the binding of recombinant Uhrf1 to indicated DNA probes. W and C indicate methylation of Watson vs Crick strands, respectively. Multiple complexes likely reflect multiple molecules of Uhrf1 bound. (G) Southern analysis of 5mC in wild-type*, dmt5Δ*, *clr4Δ, uhf1Δ* and double mutant *clr4Δ uhf1Δ* strains. (H) Whole-genome bisulfite sequencing (WGBS) analysis of wild-type, *dmt5*Δ, *clr4*Δ, *uhf1*Δ and *clr4*Δ *uhf1*Δ strains. Shown are the data for chromosome 13. (I) Number of methylated sites in the mutants analyzed in (H) as determined by WGBS. (J) Regulatory circuitry of DNA methylation in *C. neoformans*.

To understand how *C. neoformans* maintains 5mC with only a single DNMT gene, we undertook integrated biochemical, genetic, and evolutionary analyses. To our surprise, we have found that Dnmt5 is a maintenance methylase with exquisite specificity for hemimethylated DNA. Like the mammalian maintenance methylase Dnmt1, Dnmt5 is guided by H3K9me and by an orthologs of Uhrf1. These results prompted us to ask how 5mC originated in this lineage. Our phylogenetic analysis of whole genomes from Tremellaceae revealed that the ancestor of *C. neoformans* encoded for a second DNMT, which we named DnmtX. Introduction of DnmtX genes from extant species into cells that lack 5mC but contain Dnmt5 triggered cytosine methylation, indicating that DnmtX is a *de novo* methylase. Remarkably, the DnmtX gene was lost between 150-50 MYA. Thus, we investigated how 5mC has been maintained in *C. neoformans* without an efficient *de novo* system. Experimental evolution experiments demonstrate highly faithful replication of 5mC patterns. Rare, apparently random 5mC loss events were observed and substantially rarer gain events were also seen. To examine the natural evolution of 5mC, we analyzed eight isolates of *C. neoformans* from four clades separated by ∼5 MY by long read sequencing, whole genome assembly, 5mC determination, alignment, and statistical analysis. These studies revealed significant evolutionary conservation of 5mC patterns and high levels of 5mC in retrotransposons, further strongly supporting a role for natural selection. We propose that 5mC has been maintained in this lineage for millions of years without a dedicated *de novo* system by processes analogous to those that mediate the Darwinian evolution of DNA sequences: faithful replication of methylation patterns by the Dnmt5 maintenance system, variation by epimutation and natural selection for transposon silencing. We discuss the implications of the discovery of this long-lived mode of epigenome evolution.

## RESULTS

### Factors that promote 5mC in *C. neoformans*

In *C. neoformans,* regions of the genome decorated with 5mC coincide with those we reported to display H3K9me (Dumesic et al., 2015; Huff and Zilberman, 2014). Therefore, we tested whether the CD of Dnmt5 recognizes this mark. Binding of a purified fragment containing the CD to a commercial array of modified human histone peptides yielded signals with H3K9me-containing peptides as well as those containing H3K27me (Fig S1A,B). However, because H3 in *C. neoformans* differs from the human H3 sequence around lysine 27, we assessed the binding capability of the chromodomain using fluorescence polarization and peptides corresponding to *C. neoformans* H3 sequences (Fig. 1B and Fig S1C). Binding to H3K9me (K_d_=1.5 µM for H3K9me3) was substantially stronger than binding to H3K27me (K_d_>250µM), suggesting that H3K9me is the principal binding partner of the Dnmt5 chromodomain.

To assess whether H3K9me affects 5mC, we deleted the gene encoding the only H3K9 methyltransferase in *C. neoformans*, Clr4 (*clr4*Δ) (Dumesic et al., 2015). 5mC was assayed by digestion of genomic DNA with a methylation-sensitive enzyme (HpyCH4IV; A^CGT) followed by Southern hybridization. Based on published maps of 5mC in wild-type *C. neoformans*, we designed a hybridization probe directed to a unique region of centromere 13 (Fig. 1C, probe U). In addition, another probe recognizing repetitive centromeric sequences (probe R) was created to analyze the distribution of 5mC across several centromeric sites at once.

In the regions probed, 5mC levels are markedly reduced in the *clr4*Δ strain (Fig 1D) and ChIP-seq analysis of FLAG-Dnmt5 demonstrated that its recruitment to H3K9me domains is dramatically reduced (Fig. S2A). In contrast, a mutation of the Dnmt5 chromodomain that abolishes its binding to H3K9me (*W87AY90A*) has only a minor effect on 5mC, suggesting that other H3K9me-dependent factors contribute to 5mC levels (Fig. 1E and Fig. S2B). The heterochromatin protein 1 (HP1) family includes several conserved proteins that, by binding to H3K9me, participate in heterochromatin formation and in some organisms contribute to 5mC methylation (Bannister et al., 2001; Lachner et al., 2001; Lewis et al., 2010). In *C. neoformans*, we identified a single HP1 ortholog (CNAG_03458), which we named Swi6 based on its characterized ortholog in fission yeast (Fig. S2C). While the *swi6*Δ mutant marginally impacts 5mC, a *swi6*Δ *dmt5*-*W87AY90A* double mutant reduces the levels of 5mC to those seen in the *clr4*Δ strain (Fig. 1E). Immunoprecipitation-mass spectrometry and coimmunoprecipitation analysis revealed that Swi6 and Dnmt5 physically associated (Fig. S2D-E) suggesting that H3K9me promotes 5mC by recruiting Dnmt5 in two ways: via the CD of Dnmt5 and via HP1 (Fig. S2F).

The residual 5mC seen in cells lacking H3K9me led us to seek additional factors involved in DNA methylation. *C. neoformans* possesses one Uhrf1-like protein that contains an SRA domain (CNAG_00677, Fig.1F) but lacks the Tudor H3K9me reader and RING E3 ligase domains found in its human ortholog. Using native gel electrophoresis, we tested the ability of recombinant *C. neoformans* Uhrf1 to bind labelled DNA that is unmethylated, hemimethylated or symmetrically methylated at six CG dinucleotides. This experiment demonstrated that Uhrf1 selectively binds hemimethylated DNA (Fig. 1F; multiple complexes seen likely reflects multiple molecules of Uhrf1 bound), a result confirmed by competition experiments (Fig. S3). The absence of Uhrf1 (*uhf1*Δ) has almost no detectable impact on 5mC *in vivo*, but modestly alters 5mC patterns at sites probed when combined with a loss of Clr4 (*clr4Δ uhf1Δ)* (Fig. 1G).

To assay these genotypes genome-wide, we performed whole-genome bisulfite-sequencing (WGBS) of DNA extracted from wild-type, *dmt5Δ, clr4Δ, uhf1Δ* and *clr4Δ uhf1Δ* strains (Fig. 1H). The total number of symmetrically methylated CG dinucleotides, which is reduced to 31% in *clr4*Δ, drops to 11% in the *clr4Δ uhf1Δ* double mutant (Fig. 1I,J; *n.b.*: no non-CG methylation was observed). This analysis indicates that Uhrf1 and Clr4 promote 5mC through two parallel pathways (Fig 1J).

### Purified Dnmt5 displays exquisite maintenance-type specificity

*C. neoformans* Dnmt5 functions with a Uhrf1 homolog and itself recognizes H3K9me, which are features reminiscent of the mammalian maintenance methylation system. We therefore tested whether Dnmt5 has the substrate specificity expected of a maintenance enzyme. We expressed and purified full-length Dnmt5 in *S. cerevisiae*. We performed methyltransferase assays using a S-[^3^H-methyl]-adenosyl-methionine donor and synthetic 20 bp double-stranded DNA oligonucleotide substrates that harbored six CG sites. The substrate was either unmethylated, hemimethylated at all CG sites on the Watson strand, hemimethylated at all CG sites on the Crick strand or symmetrically methylated on both stands. In the absence of ATP, no activity was observed on any substrate, as compared to control reactions that lacked Dnmt5 (Fig. 2A). However, in the presence of ATP, methyltransferase activity was observed only on hemimethylated substrates with apparent catalytic rates comparable to that of human Dnmt1 (Fig. 2A, B)(Pradhan et al., 1999). Endpoint assays with the unmethylated substrate did not reveal any activity above background on unmodified DNA; under the same conditions, we observed methyltransferase activity on the hemimethylated substrate that was ∼1300-fold over background. (Fig. S4B). In addition, Dnmt5 activity is specific for CG dinucleotides since no activity was detected on DNA substrates containing CHG motifs instead of CG motifs, regardless of cytosine methylation state (Fig. S4C).

**Figure 2.**
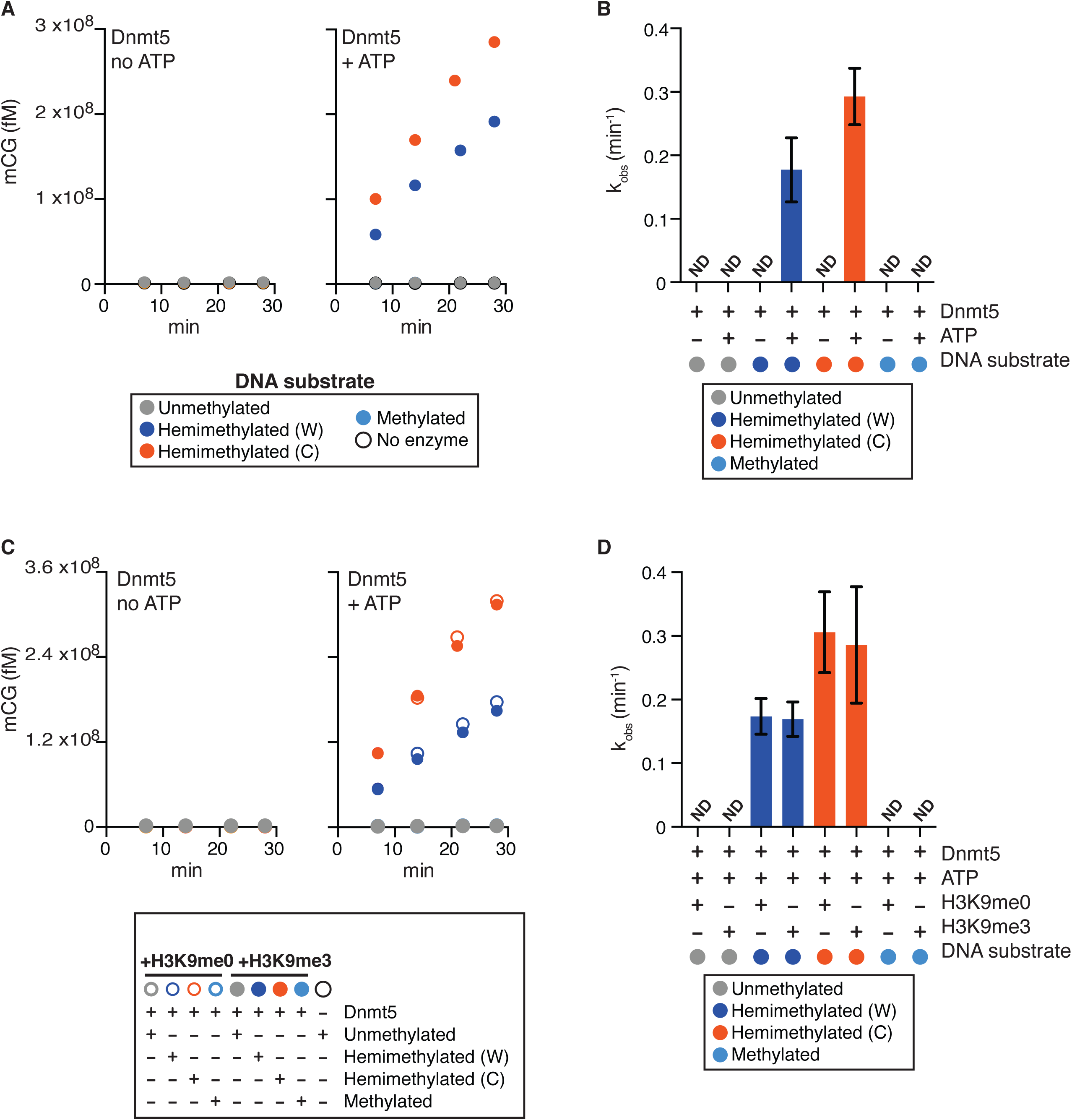
Hemimethylated DNA but not unmethylated DNA is a substrate of purified Dnmt5. (A) Reaction curves showing incorporation of label from S-[methyl-^3^H]-adenosyl-L-methionine into the indicated DNA oligonucleotide substrates using 30 nM Dnmt5 and 5 μM DNA substrate, in the presence or absence of 1 mM ATP. (B) Initial rates of Dnmt5 activity on DNA substrates described in (A). n=3-4; error bar represents SD. ND: not detected. (C) Reaction curves of incorporation of label from S-[methyl-^3^H]-adenosyl-L-methionine into the indicated DNA oligonucleotide substrates using 30 nM Dnmt5 and 5 μM DNA. Reactions were performed in the presence or absence of H3K9me0/3 peptide (5 μM) and ATP (1 mM). (D) Initial rates of Dnmt5 activity on DNA substrates described in (C). n=3-4; error bars represent SD. ND: not detected.

Because the chromodomain of Dnmt5 recognizes H3K9me, we asked whether *de novo* activity could be detected when trimethylated H3K9 peptide was added to the *in vitro* reaction. Rates on hemimethylated substrates were comparable in the presence of methylated or unmethylated H3K9 peptide, and no activity was detected on unmethylated DNA (Fig. 2C and D). We also tested whether recombinant Uhrf1 and Swi6 could trigger a *de novo* activity when added to the reactions, but none could be detected (Fig. S4D). We conclude that Dnmt5 is a maintenance enzyme *in vitro* with activity on hemimethylated substrates and no detectable activity on unmethylated substrates under a range of conditions.

### 5mC is propagated by a maintenance mechanism *in vivo*

Although Dnmt5 displays the *in vitro* substrate specificity expected for a maintenance type enzyme, it may function *in vivo* with cofactors that could change its properties. A way to address this question is to remove Dnmt5 from a cell, allow methylation to be lost, and then re-introduce the enzyme. If Dnmt5 behaves as a maintenance enzyme in cells, DNA methylation should not be globally restored since its hemimethylated substrate would be lost.

We placed the gene for Dnmt5 under the control of a galactose-inducible promoter (Fig. 3A; *pGAL-DMT5*). This allele was tagged at its N-terminus with a 2XFLAG epitope tag. In the first series of experiments, the corresponding targeting construct was introduced by transformation and recipient cells were selected under conditions repressive for the *pGAL7* promoter (*R*). After induction of the Dnmt5 protein using the inducer galactose for 20 (I_20_) or 45 generations (I45), we were not able to detect 5mC by Southern hybridization, although expression of the construct was evident (Fig. 3A). To assess if the *pGAL-DMT5* allele is functional, the targeting construct was transformed into wild-type cells but this time recipient cells were selected under conditions that induce pGAL7 (Fig. 3B). The 5mC pattern of the strain under these inducing conditions (I_0_) is indistinguishable from that of wild-type indicating that the inducible, tagged allele is indeed functional. Subsequently, upon repression of *pGAL-DMT5*, 5mC was lost. However, again, re-induction of Dnmt5 for 40 (*I_40_*) or 90 generations (I_90_) did not restore detectable 5mC. To determine whether 5mC could be restored if it was only partially depleted from cells, for example by a nucleation-spread mechanism, we set up a time-course experiment in which we repressed *pGAL-DMT5* for limited number of generations (*R*) and re-induced (*I*) it. In repressing conditions, substantial decrease of 5mC signal was observed only after 25 generations, which correlated with the depletion of the mRNA produced by the p*GAL-DMT5* overexpression construct (Fig. S5). However, upon transfer to galactose-containing inducing medium, we did not observe restoration of 5mC (Fig. 3C; corresponding quantification shown in Fig. 3D).

**Figure 3.**
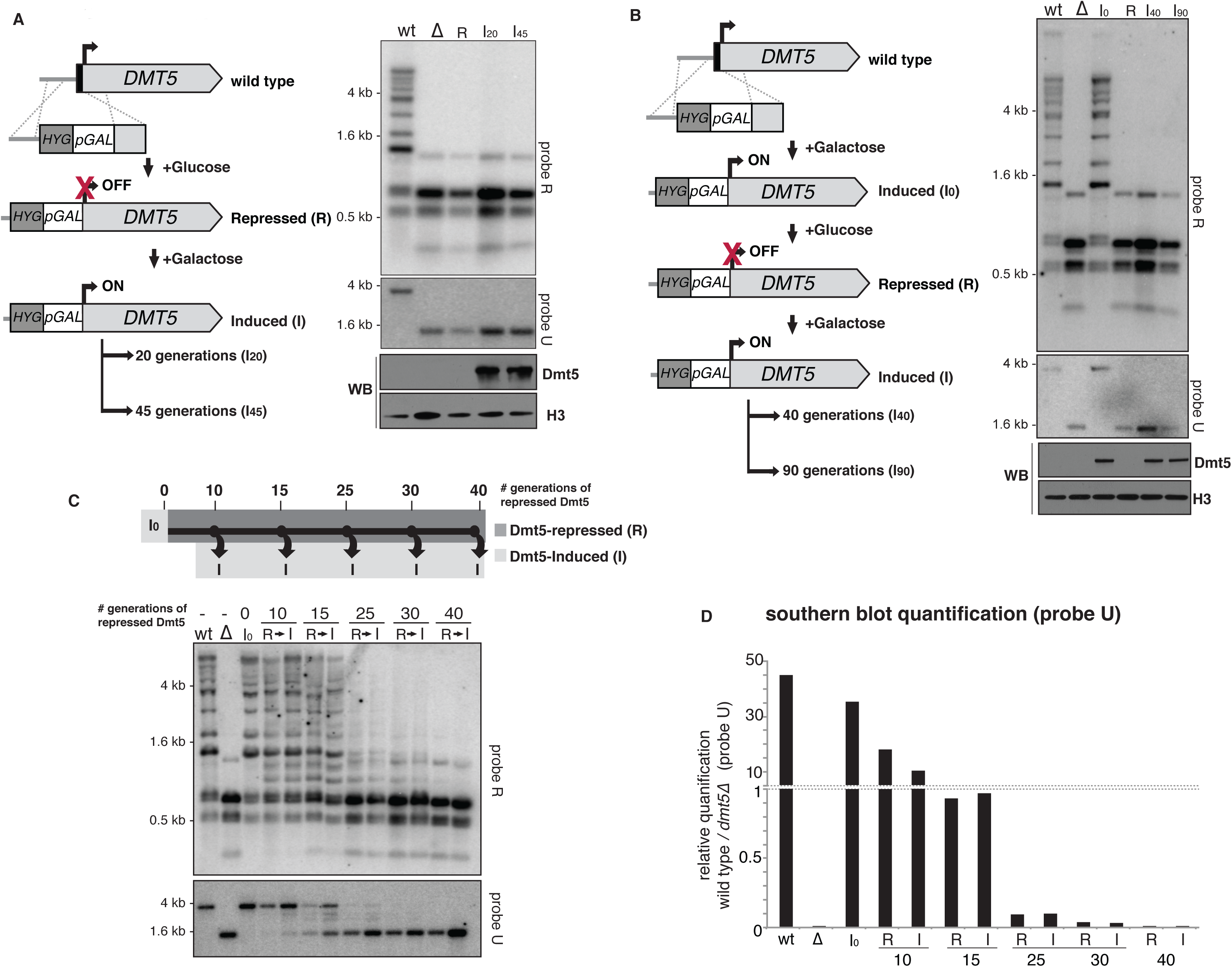
Transient repression of Dnmt5 results in sustained loss of 5mC. (A) OFF-ON scheme. Knock-in of a galactose-regulated *GAL7* promoter and epitope tag upstream of the Dnmt5 coding sequence was performed as shown. Strains were selected under repressing (glucose) conditions (R). Dnmt5 was induced by addition of galactose for 20 (I_20_) or 45 generations (I_40_). Strains were analyzed by Southern hybridization as in Figure 1. WB: Western blot indicating levels of FLAG-Dnmt5 and histone H3. (B) ON-OFF-ON scheme. Knock-in of a galactose-regulated *GAL7* promoter and FLAG epitope-tag upstream of the Dnmt5 coding sequence was performed as shown. In contrast to (A), selection for the knock-in allele was carried in presence of galactose (Induced-I_0_). Dnmt5 was then repressed to produce a loss of DNA methylation (R) and subsequently induced for 40 (I_40_) or 90 generations (I_90_). Strains were analyzed by Southern hybridization. (C) ON-OFF-ON scheme with variable OFF times. Strain produced under inducing conditions as in (B) was subject to the indicated number of generations of repression (R) followed by 20 generations of induction. Southern analysis was performed as in (A). (D) Quantification of the ratios of the band at 3.5 kb (wild-type) divided by the band at 1.5 kb (*dmt5Δ)* for probe U calculated for each sample in (C) using ImageJ software.

We performed an analogous analysis using a different approach. We first deleted a portion of the *DMT5* gene followed by the re-introduction of the missing DNA sequence (Fig. 4; the re-introduced allele is termed *RI-DMT5;* see materials and methods). Although the expression of *RI-DMT5* was comparable to that of a wild type allele, we were not able to detect 5mC by Southern hybridization (Fig. 4A). WGBS sequencing of the *dmt5*Δ strain revealed no methylation on any CG sites genome wide (Fig 4B). Analysis of two independently-derived *RI-DMT5* strains revealed that methylation is also globally absent save for two sites detected in each strain (Fig. 4B; Fig. S6A). To examine methylation using an independent global method, we employed attomole-sensitivity mass spectrometry to measure 5mC. Analysis of 5mC levels present in gDNA extracted from wild type, *dmt5Δ* and *RI-DMT5,* revealed 0.33% 5mC in wild type, none detectable in DNA extracted from the *dmt5*Δ strain and trace amounts (detectable but below quantification limits) in DNA extracted from the *RI-DMT5* strain (Fig 4C). This trace activity suggests that Dnmt5 may able to act at a very low level on unmodified DNA, an event which would be then ‘remembered’ by its maintenance activity. Investigations of methylation dynamics in wild type cells are described further below.

**Figure 4.**
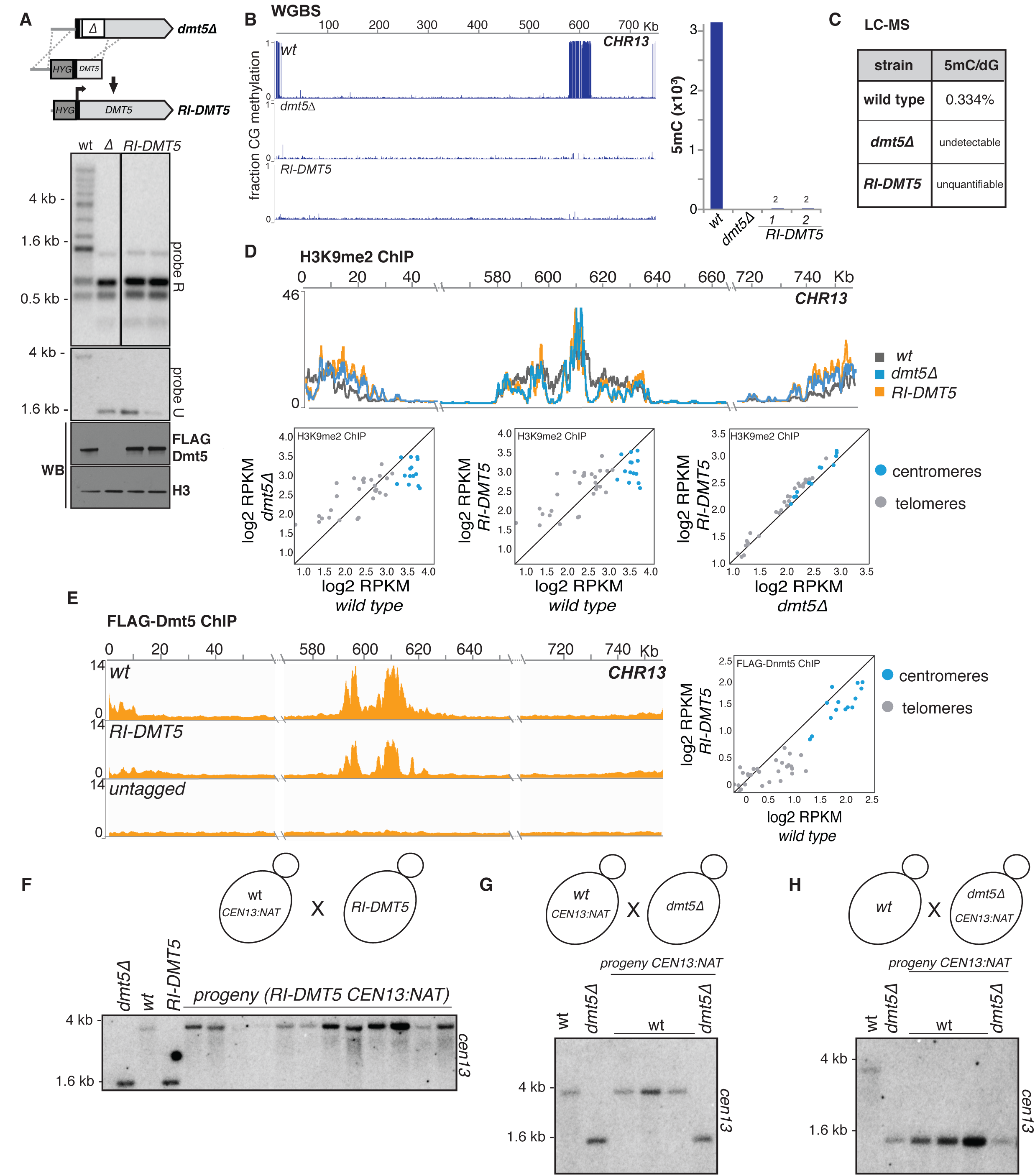
Genetic loss of DNA methylation is mitotically and meiotically irreversible. (A) Re-introduction (RI) scheme. A partial knockout of the *DMT5* locus was produced and then corrected by homologous recombination and N-terminally tagged as in Figure 2 to produce the RI-DNMT5 strain. A hygromycin resistance marker (*hygR*) is inserted upstream of the Dmt5 gene during this process. DNA methylation was assessed as in Figure 1. (B) WGBS analysis of *RI-DMT5* strain. Shown are data for chromosome 13 for the indicated genotypes. Bar graph shows number of called symmetrically methylated CpGs for the indicated genotypes. (C) Quantification through mass spectrometry of 5mC in DNA of wild type, *dmt5*Δ and *RI-DMT5* strains (D) Analysis of H3K9me2 signals in wild type, *dmt5*Δ and *RI-DMT5* strain. ChIP-seq signal for H3K9 methylation shown as normalised to signal for the H3 ChIP. Shown are the data for centromere 13 for the indicated genotypes (top) and the pairwise comparison of the total RPKM counts for each centromere (blue) and telomere (grey) (scatter plot, bottom). (E) Analysis of Dnmt5 recruitment in *the RI-DMT5* strain. ChIP-seq signal for FLAG-Dnmt5 were normalised to signal for the input samples. Shown are the data for centromere 13 for the wild type, *RI-DMT5* and a strain not expressing FLAG-tagged proteins. (F) Genetic cross testing the functionality of RI-Dnmt5 protein. Centromere 13 of a wild-type strain was tagged by insertion of a nourseothrycin resistance gene 3.5 Kb from its left end (*CEN13*::*natR)* and the resulting strain was crossed to a strain harboring the *hygR*-marked *RI*-*dnmt5* allele. Progeny strains doubly resistant to hygromycin and nourseothrycin were selected. Strains were analyzed for as described in Figure 1 using probe U, which is specific for chromosome 13. (G) Control cross. Dnmt5-harboring cells containing *CEN13::natR* were crossed to *dnmt5Δ* cells. Progeny harbouring the Dnmt5 and *CEN13::natR* were analyzed as in (C). (H) Experimental cross testing whether sexual reproduction enables de novo methylation. *dmt5*Δ cells harboring *CEN13::natR* were crossed to wild-type cells. Progeny harbouring a wild-type Dnmt5 gene and *CEN13::natR* were analyzed as in (C).

We observed that H3K9me is still globally maintained in both the *dmt5*Δ strain as well as in the *RI-DMT5* strain, although the distribution is altered compared to wild-type depending on the genomic region with an increase of the signal at subtelomeric regions (Fig. 4D, Fig. S7A). Furthermore, ChIP-seq experiments demonstrate that the *RI-DMT5* allele still localizes to heterochromatic regions (Fig. 4E; Fig. S7B).

To test explicitly if *RI-DMT5* is capable of efficiently maintaining DNA methylation, a wild-type strain whose centromere 13 was marked with a linked drug resistance marker (*CEN13::natR*) and crossed to the *RI*-*DMT5* (Fig. 4F). From the cross, meiotic progeny bearing the *RI-DMT5* allele and the marked centromere were selected. 5mC was assayed by Southern hybridization using a probe specific for *CEN13*. All progeny tested displayed a wild-type 5mC pattern (Fig. 4F), indicating that the *RI-DMT5* allele is functional when provided with a methylated substrate.

In many species, sexual reproduction is required for the establishment of 5mC by one or more *de novo* enzymes. It has been reported that a tandemly-repeated *URA5*-marked transgene can be heritably silenced after mating and meiosis/sporulation in *C. neoformans*, but that this silencing occurs via RNAi and not via DNA methylation(Wang et al., 2010). Nonetheless, to determine whether an unmethylated centromere can be methylated during sexual reproduction, we performed two genetic crosses and analyzed the meiotic progeny. In a control cross, we crossed a *CEN13::natR* strain with a *dmt5*Δ strain and analyzed three progeny that contained both the marked *CEN13* and wild-type *DMT5* allele by Southern hybridization using a probe specific for *CEN13*. As expected, 5mC was maintained at the methylated centromere (Fig. 4G). In the experimental cross, we crossed a wild-type strain to one harboring a *dmt5*Δ allele that also carries *CEN13::natR.* We again analyzed three progeny that express the wild-type *DMT5* gene and the marked centromere, which entered the cross in an unmethylated state. We observed that such progeny remained unmethylated, indicating that Dnmt5 is able to maintain methylated DNA but does not re-establish on a centromere that had completely lost methylation (Fig. 4H). Thus, sexual reproduction does not lead to the efficient restoration of 5mC. Finally, we examined cells exposed to several stress conditions to determine whether they could reestablish methylation in the *RI-DMT5* strains. We did not observe restoration of methylation (Fig. S8).

To assess whether Dnmt5 is able to maintain DNA methylation independently on non-endogenous sequences, we methylated *in vitro* a targeting construct containing a DNA fragment rich in HpaII sites with the HpaII methyltransferase (which produces symmetric CmCGG) and integrated it into the *C. neoformans* genome at the right border of centromere 4 in order to place it in proximity to H3K9me-modified chromatin (Fig. 5A-C). DNA from the transformants was digested with the 5mC-sensitive HpaII restriction endonuclease and qPCR was performed using a primer pair that spanned the cleavage sites to measure the fraction of a particular site that was methylated (Fig. 5D). Analysis across seven different HpaII sites revealed that all were maintained *in vivo* (Fig. 5D). Thus, there appears to be no obligate sequence requirements beyond CG dinucleotides for methylation to be maintained *in vivo*, a finding consistent with our biochemical results.

**Figure 5.**
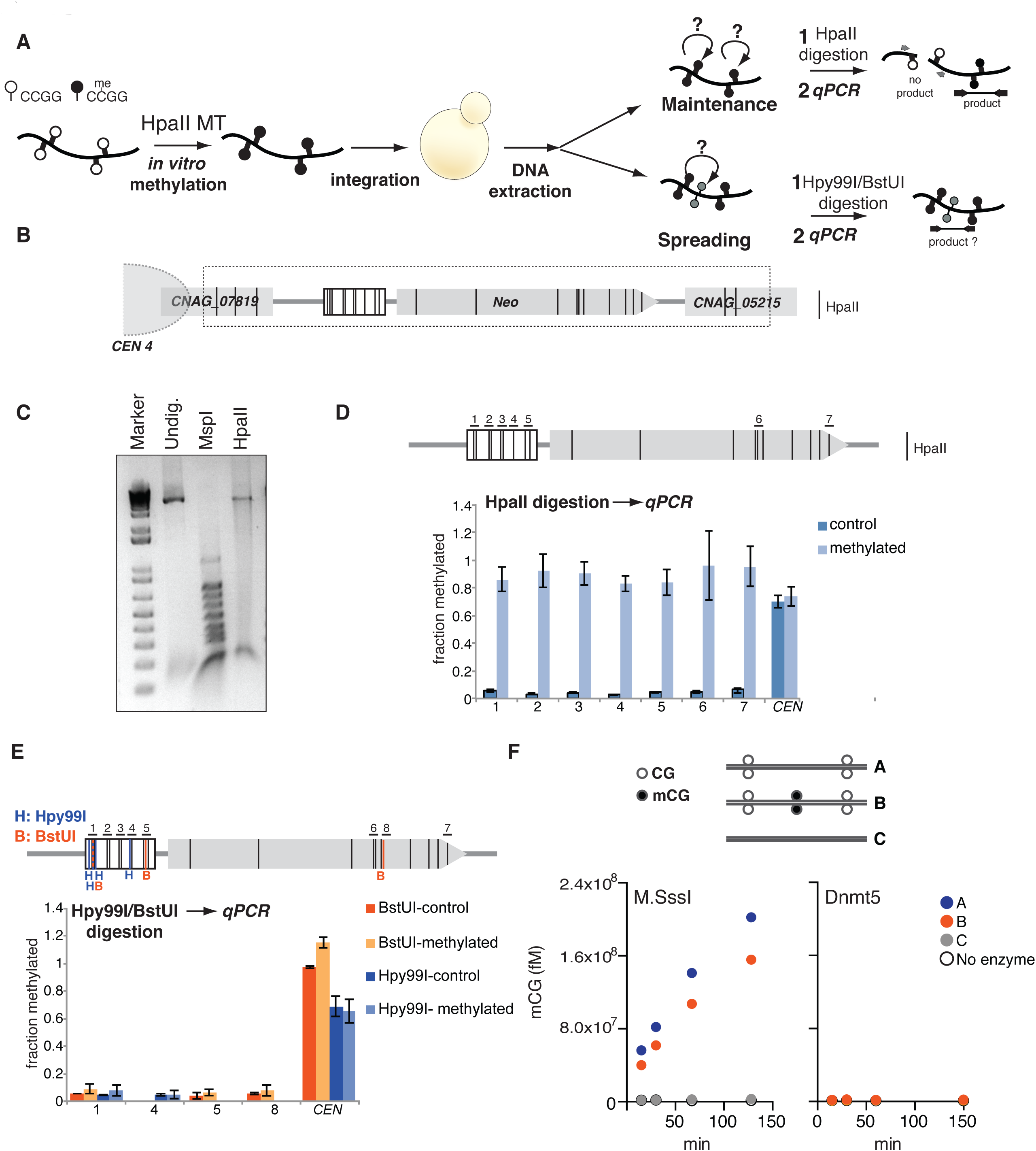
Dnmt5 maintains *in vitro* methylated DNA. (A) A DNA fragment was methylated *in vitro* using the HpaII DNA methyltransferase (HpaII MT) and subsequently integrated into *C. neoformans*. gDNA extracted from isolates that correctly integrated the fragment, was digested with the restriction enzyme HpaII. To test the maintenance of the *in vitro*-methylated 5mC, qPCR was performed on region spanning the restriction sites. To test the ability of Dnmt5 to spread DNA methylation, gDNA was digested with either Hpy99 or BstUI and qPCR was performed across these restriction sites. White and black lollipops represent unmethylated and methylated sites, respectively. (B) Schematic representation of the construct (shown within the dashed box) and its integration adjacent to centromere 4 (*CEN4*). Vertical black lines represent the site (CCGG) recognised by the HpaII methylatransferase. (C) Agarose gel of undigested *in vitro* methylated DNA (Undig.) digested with the methylation insensitive restriction enzyme MspI (MspI) or its methylation-sensitive isoschizomer HpaII (HpaII). (M: Marker). (D) qPCR performed on gDNA extracted from yeast transformed with the control unmethylated (control, dark blue) and *in vitro* methylated construct (methylated, light blue). gDNA was digested with HpaII and qPCR was performed as described in A on regions 1-7. qPCR was normalised over a region of the genome that is not cleaved by HpaII. *CEN*: positive control-region on centromere 13 that contains a methylated HpaII site. Black vertical lines indicate site of HpaII cleavage. (n=3, error bars represented as SD). (E) gDNA digested with either the restriction enzymes Hpy99I or BstUI. qPCR was performed on gDNA containing the unmethylated control (dark blue, Hpy99I-control and orange, BstUI-control) or the *in vitro* methylated construct (light blue, Hpy99I-methylated and yellow, BstUI-methylated). qPCR was normalised over a region of the genome that is not cleaved by Hpy99I or BstUI. In the scheme (top), black vertical lines indicates site of HpaII cleavage, blue vertical lines indicate sites of Hpy99I and orange vertical lines indicate the sites for BstUI. (n=3, error bars represented as SD). (F) Double-stranded DNA containing two unmethylated CG (A) or unmethylated CG surrounding a central methylated CG site (B) or without any CG dinucleotide (C) were used as substrate of DNA methylation kinetics with M.SssI (left) or 30 nM Dnmt5 (right).

We tested next whether 5mC could spread *in vivo* by using the *in vitro* methylated sites as nucleation sites. We digested the genomic DNA with the 5mC-sensitive enzymes BstUI (CGCG) and Hpy99I (CGWCG) and performed qPCR across these sites. We did not observe a signal above background, indicating no detectable spread of 5mC to these nearby CG dinucleotides (Fig. 5E). We obtained similar results *in vitro*: we incubated purified Dnmt5 with a dsDNA substrate containing either no CG sites, or two unmethylated CG sites, or two ummethylated CG sites accompanied by a symmetrically methylated CG site (Fig. 5F). Although methylation products were detected upon incubation with a control bacterial DMT M.SssI, no signal was detected upon incubation with Dnmt5, indicating that the enzyme does not display detectable *de novo* activity even when presented with a nearby methylated site (Fig. 5F).

### The gene for a *de novo* methylase was lost in an ancestral species

Given that Dnmt5 is the sole DNMT predicted by the *C. neoformans* genome, and seems to behave as a maintenance DNMT, how was 5mC established? To approach this question, we examined the genomes in the family of organisms to which *Cryptococcus neoformans* belongs, the *Tremellaceae* (Fig. 6). The genomes of species closely related to *C. neoformans*, including the human pathogen *Cryptococcus gattii* and the nonpathogens *Cryptococcus amylolentus* and *Cryptococcus wingfieldii*, harbor a single predicted DNA methyltransferase, which is orthologous to Dnmt5 (Fig. 6A). The exception is *C. deuterogattii* which has inactivated all transposons and lost both RNAi and 5mC (Yadav et al., 2018b). The next closest species, *Cryptococcus depauperatus*, lacks predicted DNA methyltransferases, also indicating loss of *DMT5* during its evolution (Fig. 6A). Strikingly, more distant species encode both Dnmt5 and an uncharacterized predicted DNA methyltransferase, which we term DnmtX (genetic locus: *DMX1*; Fig. 6A). This predicted protein contains a bromo-associated homology (BAH) domain and a Dnmt catalytic domain. This phylogenetic pattern indicates that the *DMX1* gene was lost in the common ancestor to *C. neoformans* and *C. depauperatus* and that *C. depauperatus* subsequently lost the *DMT5* gene. Given the fact that Dnmt5 is a maintenance methylase, we hypothesized that DnmtX is a *de novo* methyltransferase.

**Figure 6.**
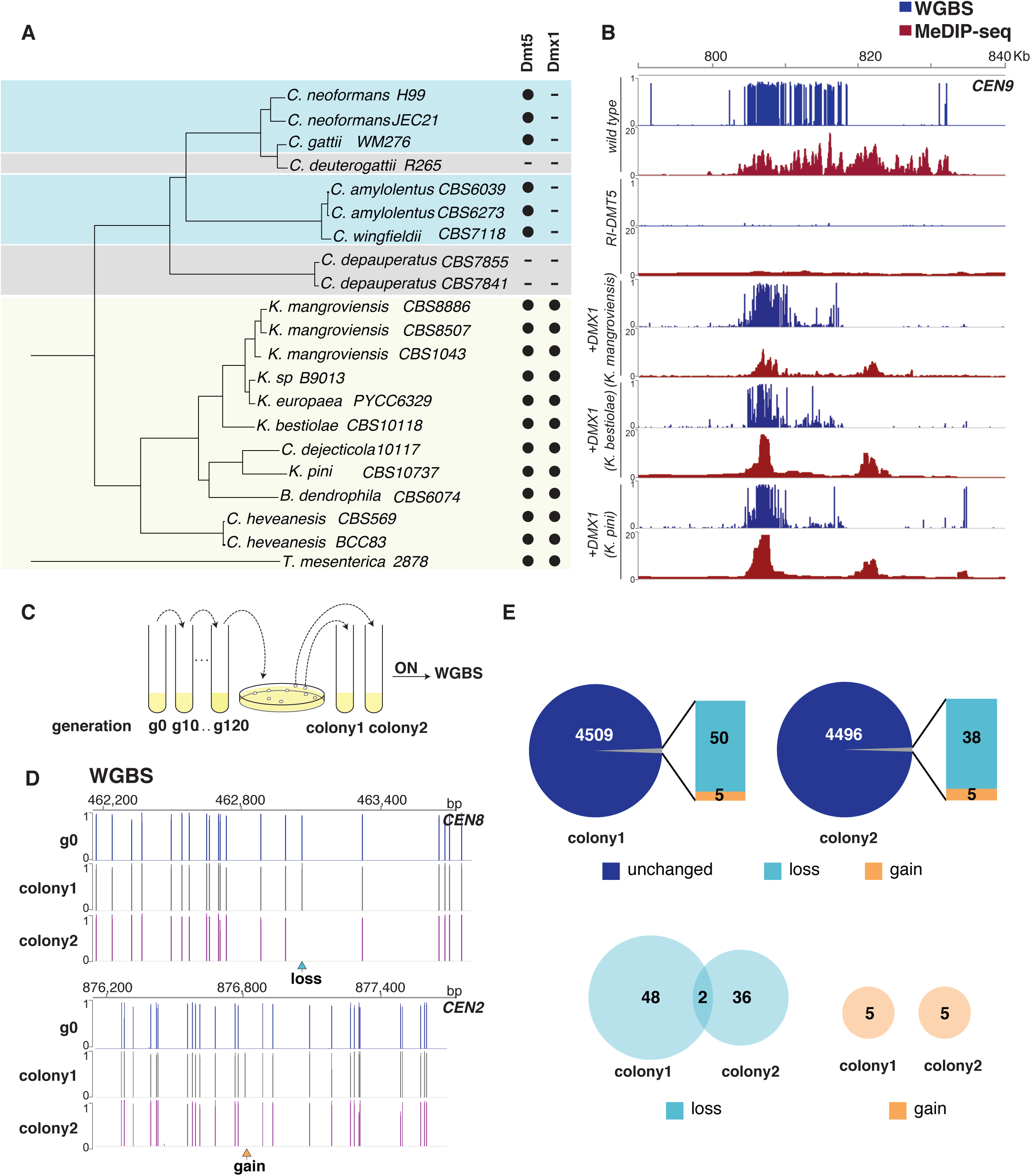
Ancient loss of gene coding for a *de novo* Dnmt in an ancestor of *C. neoformans* and laboratory evolution experiments. (A) Phylogenetic analysis of whole-genome sequences. Shown are the phylogenetic relationships between *C. neoformans* and characterized *sensu stricto* and *sensu lato* groups. Phylogeny is based on the analysis of whole-genome sequences (C. Cuomo et al., manuscript in preparation). The genomic presence of gene coding for orthologs of Dnmt5 and of a second DNMT (DnmtX) are indicated by filled circles. As indicated by the arrows, gene for DnmtX was lost prior to the divergence of *C. neoformans* and *F. depauperatus*, and the gene for Dnmt5 was subsequently lost in the *F. depauperatus* lineage as well as in the *C. deuterogattii* lineage. (B) Expression of extant DnmtX-encoding genes in an *RI-DMT5 C. neoformans* strain triggers *de novo* 5mC. Three DnmtX from indicated organisms were expressed in a RI-DNMT5 strain. 5mC was assessed by MeDIP-seq (red) and WGBS (blue). MeDIP signal is shown as ratio over *dmt5Δ.* Shown are data for centromere 9. (C) Scheme for laboratory evolution assay. Wild-type cells (g0: generation 0) were propagated in rich media and for 120 generations and then plated on agar plates to form single colonies. Two independent isolates (colony 1 and colony 2) were grown overnight in rich media and the gDNA extracted processed in technical duplicates for BS-seq. (D) WGBS of a portion of the starting culture (g0) and the two different isolates grown for >120 generations (colony 1 and colony 2). Shown are the reproducible sites between technical replicates on centromere 9. Sites that are present in wild-type and absent after >120 generation are marked as “loss” while sites absent in wild-type but occurring in one of the isolates are marked as “gain”. (E) Differential methylation analysis (DMS) was performed to compare g0 with the propagated isolates. The majority of reproducible sites remained unchanged (dark blue) and only a small fraction showed variation compared to g0. Specifically, in light blue is shown the number of sites that have been lost and in orange are shown novel sites arising after propagation. (F) Number and relationship of shared lost (left) and gained (right) sites between colony 1 and 2.

To test this hypothesis, we cloned the genes for DnmtX from *Kwoniella mangroviensis, Kwoniella bestiolae* and *Kwoniella pini*. Each of these was placed under the control of the *pGAL7* galactose-inducible promoter, tagged at the N-terminus with a hemagglutinin epitope tag, and then introduced as a transgene into a *C. neoformans* strain in which the gene for Dnmt5 was disrupted and then restored with a FLAG-tagged allele (*RI-DMT*5). We chose this approach because we predicted that establishment of 5mC by a DnmtX enzyme might be inefficient, especially if cofactors important for *de novo* methylation were not also introduced.

We induced expression of DnmtX in each strain using medium containing galactose (Fig. S9) and DNA was then extracted. The levels of 5mC were assessed genome-wide by two technically orthogonal methods, methylated DNA immunoprecipitation followed by high-throughput sequencing (MeDIP-seq) and WGBS. In both assays, we observed the broad accumulation of 5mC in each of the three strains, primarily at centromeric regions (Fig. 6B, Fig. S10 and S11). These data indicate that all three DnmtXs act as *de novo* methylases *in vivo*, providing strong evidence that the ancestral DnmtX also had this function.

### Experimental evolution reveals fidelity of 5mC maintenance

The loss of the ancestral DnmtX gene occurred prior to the divergence of *C. neoformans* and *C. depaueratus* but after the divergence of *Tremella mesenterica* and *Kwoniella hevanensis* (Fig. 6A). The divergence time for *C. neoformans* and *C. gattii* is estimated to be between 34 and 49 MYA (D’Souza et al., 2011; Ngamskulrungroj et al., 2009). Given the phylogenetic relationships, the divergence time of *C. neoformans* and *C. depauperatus* would be considerably more ancient. It has been estimated that the common ancestor of *C. neoformans* and *T. mesenterica* lived 153 MYA (Floudas et al., 2012). Thus the *DMX1* gene was likely to have been lost between roughly 150 and 50 MYA. Taken together, these data indicate that 5mC has been maintained by Dnmt5 over a remarkably long period of time in the population that gave rise to the pathogenic *Cryptococcus* species complex. In principle, two non-mutually exclusive models could promote 5mC in *C. neoformans*. At one extreme, 5mC has been maintained since the loss of DnmtX by a combination of the fidelity of inheritance and natural selection. Alternatively, rare *de novo* events (as seen above) might occur with sufficient frequency that, together with selection, maintain an overall level of methylation at the centromere.

To test the plausibility of these models, we performed an experimental evolution experiment. A single colony from wild type cells (generation 0, or g0) was grown in liquid rich media for 120 generations without bottlenecking the population. Dilutions of the final culture were plated to form single colonies on agar plates. Two independent clones were selected and replicate WGBS analysis was performed on their genomic DNA as well as DNA extracted from a portion of the parental strain (Fig. 6C). We detected 50 and 38 loss events, respectively, (Fig. 6D, E), indicating that 99.0% and 99.2% of 5mC sites were maintained after 120 generations. Only two sites were lost in both colonies, an overlap expected by chance (Fig. 6E, Table S1). Assuming all sites are equally maintained, this would correspond to a per-generation loss rate of 8.4 x 10^-5^. Consistent with the reintroduction experiments described above, we observed five gain events in each experiment (nonoverlapping; Fig. 6E, Table S1). Because these gain events are considerably more infrequent than the loss events, the system is not at equilibrium under laboratory conditions. Thus, selection on methylation levels (across centromeres or at specific sites) likely operates in natural populations to maintain the steady state of methylation. These experimental evolution data are consistent with both models described above: a high fidelity of maintenance coupled with selection is consistent with a long-term maintenance model, while our detection of rare gains indicates that gain-type epimutations are also injected into the system.

### Evolutionary analysis of 5mC patterns reveals conservation

To investigate further the inheritance of 5mC patterns, we examined its evolutionary conservation. To accomplish this, we first sought to align centromeric regions from isolates that had diverged by a significant period of time. *C. neoformans var grubii*, the species of the reference H99 strain, has four well-characterized clades, VNI, VNII, VNBI and VNBII, which shared a common ancestor approximately 4.7 MYA (Ngamskulrungroj et al., 2009) (Fig. 7A). Short-read sequences are available for several hundred strains (Desjardins et al., 2017), however, centromeric regions, where the bulk of 5mC resides, are not assembled as they contain repetitive sequences, including six families of *gypsy* and *copia* retrotransposons, Tcn1-6 (Janbon et al., 2014). Therefore, we used long read Oxford Nanopore (ONT) sequencing to assemble near-complete genomes of two isolates from each of the four clades (Fig. 7B). Dot plot analysis of centromeres revealed widespread relative rearrangements of sequence (insertions, deletion, inversions), precluding end-to-end sequence alignment (Fig. S12A). We instead adopted a local alignment approach to obtain a series of pairwise alignments between segments of centromeres of the eight assembled genomes. On average, 88.5% of centromeric sequence from a given strain could be aligned to the homologous centromere of at least one other strain, with a median alignment length of 607 bp (Fig. S12B, C). We performed replicate WGBS analysis on each strain and overlaid these data onto the alignments. An example of this analysis is shown in Figure 7C, which displays the 5mC status of CG sites that can be aligned between a centromere of an arbitrarily chosen strain (A1_35_8) and the homologous centromere on other seven strains. Next, for each centromeric CG dinucleotide in each of the strains, we determined the number of strains (N) to which a given CG nucleotide could be identified by alignment and then calculated the fraction of CGs that were methylated in all N strains (Fig. 7D). Statistical analysis revealed significant conservation of methylated sites regardless of the strain used as a reference and the number of strains to which a centromeric CG dinucleotide could be aligned.

**Figure 7.**
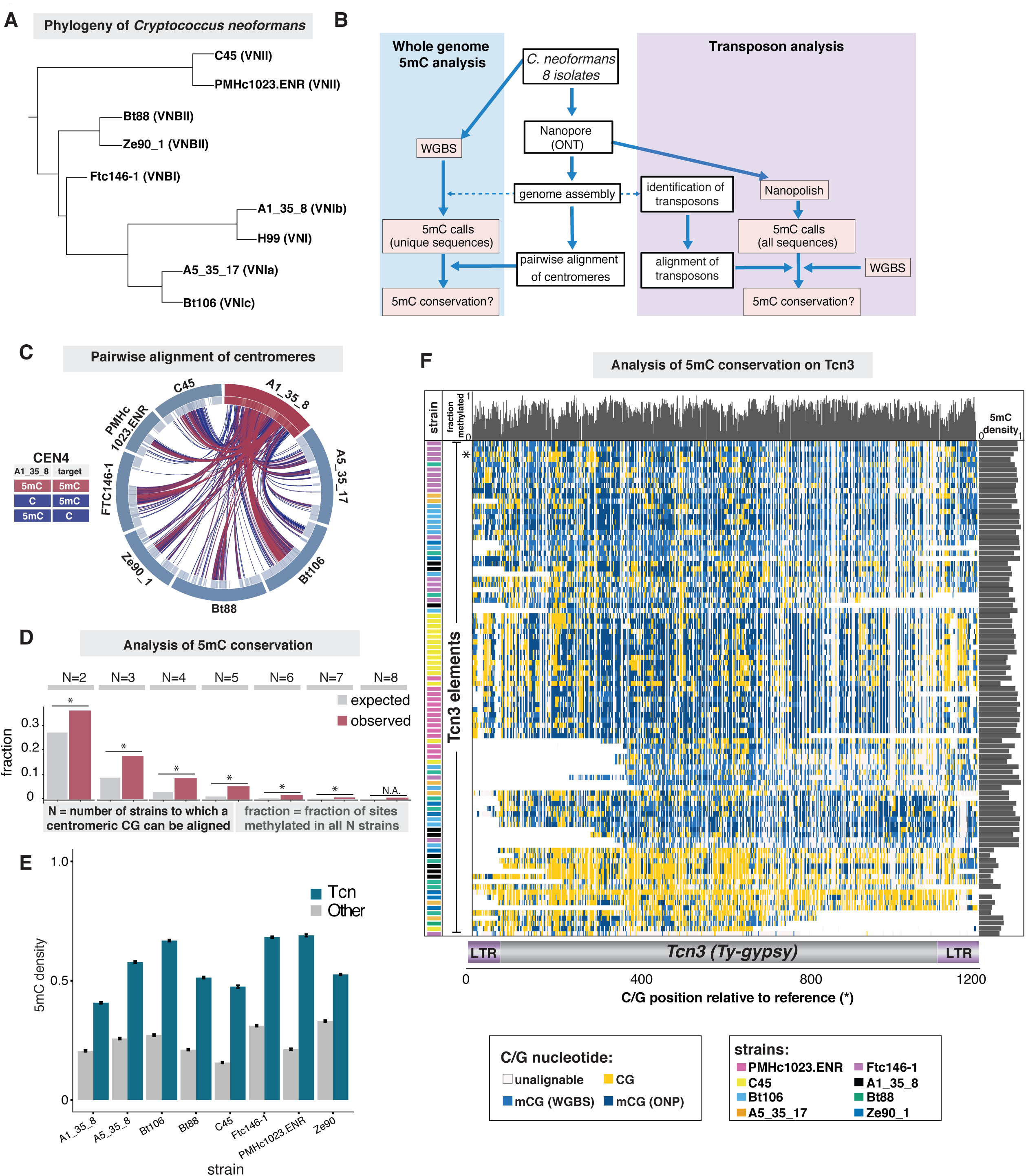
Cytosine methylation sites are conserved between isolates of the *C. neoformans*. (A) Phylogenetic relationships between isolated of *C. neoformans*. Phylogeny is based on the analysis of whole-genome sequences. (B) Pipeline for the analysis on the evolutionary conservation of 5mC. The genomes of the eight isolates related to *C. neoformans* H99 were assembled using long read sequencing and their centromeric sequences were aligned pairwise. Whole genome 5mC evolutionary analysis was performed using WGBS aligned to the assembled genomes. The 5mC data were superimposed to the sequence alignment and conservation of 5mC assessed. Transposon analysis was performed on the identified transposable elements. Nanopolish methylation calls were used to complement the 5mC calls on repetitive sequences that were absent in the WGBS analysis. Transposable elements were aligned and 5mC conservation assessed. (C) Circos plot representing the pairwise alignment of *CEN4* in A1_35_8 (red segment) with the centromeres of the other seven isolates (blue segments). The red lines represent CG dinucleotides that are methylated in both A1_35_8 and in target centromeres. The blue lines correspond to CG sites that lost 5mC in either A1_35_8 or the compared centromere. Light grey shadings represent the regions that are both alignable and display coverage in WGBS data. (D) 5mC conservation analysis. Each CG dinucleotide was aligned and sorted into seven different categories (N) according to the number of strains in which the CG could be aligned (N=2 to 8; N includes the reference strain). The fraction of CGs in which all N alignable CGs are methylated (red) is higher than that expected by chance (grey) (* p < 1×10^-17^). (E) Comparison of the mean methylation levels on centromeric sequence containing Tcn elements (in blue) and other centromeric sequences (grey) from different isolates of *C. neoformans*. (F) Alignment of 5mC in Tcn3 from the different isolates. Longest (top fifth) Tcn3 sequences were aligned to a reference Tcn3 from the strain Ftc146_1 (indicated with *) and clustered according to their 5mC status. On the bottom, schematic representation of Tcn3 (grey box) flanked by two long terminal repeats (LTR-purple boxes). On the x-axis, each box represents a C/G position relative to the reference. A white space indicates that the sequence could not be aligned; a cream box indicates C/G nucleotides was not conserved; a yellow box indicates that CG is not methylated; in light blue box the CG dinucleotides is methylated (data from WGBS analysis) and in dark blue boxes are the 5mC from Nanopore sequencing analysis. The fraction of CGs methylated for each position is display across the top (mean: 0.7039, p < 1×10^-16^, test of independence with null generated by permutations). The fraction of CGs methylated for each Tcn3 is shown on the right histogram (mean: 0.7113).

Given the concomitant loss of 5mC and intact transposons in *C. deuterogattii*, it seems likely that important targets of DNA methylation are the Tcn elements described above. We tested whether these elements display higher levels of 5mC than non-Tcn centromeric sequences and whether these patterns are conserved. In all eight strains, we identified sequences at centromeres that are highly homologous to Tcn1-6, including a fraction harboring two LTRs, which are candidates for active transposons (Fig. S12D). Due to their length and repetitiveness, Tcn elements are not well-covered by WGBS; hence we supplemented the WGBS data using methylation calls from ONT long-read data, which we found to be highly correlated with those produced by WGBS (Fig. S12E, F). Consistent with selection acting on 5mC in Tcn elements, we observed significantly more 5mC on Tcn-related sequences than non-Tcn sequences within centromeres (Fig. 7E).

Since Tcn3-related sequences displayed the longest median length and the most LTR-flanked elements (Fig. S12D), we performed additional analysis on these elements. We aligned each CG dinucleotide from the longest (top quartile) Tcn3-related sequences and overlaid the 5mC data. Strikingly, for the vast majority of CG dinucleotides aligned (Fig 7F, vertical columns), we observed high conservation of methylation (mean fraction = 0.7, p < 10^-16^) compared to a control model for Tcn3 of independently permutated 5mC sites (mean fraction = 0.4149) indicating sharing of 5mC patterns across strains above expectation. Note that in several cases we observed similar patterns of 5mC on different Tcn3 elements within a single strain (Fig. 7F); this is more likely the result of recombination events such as unequal crossing over rather than retrotransposition as the latter would be expected to erase methylation patterns.

## DISCUSSION

### A maintenance DNA methylation system in the fungal kingdom

Cytosine methylation of DNA is a widespread modification of DNA that plays numerous critical roles, particularly in genome defense, yet has been lost many times in diverse eukaryotic lineages. In animals and land plants, *de novo* enzymes establish methylation at specific developmental windows and maintenance enzymes propagate this initial activity after DNA replication. The phenomenon of methylation-induced-premeiotically (MIP) in the ascomycete *A. immersus* (Rossignol and Faugeron, 1995) suggests that the *de novo*-maintenance paradigm might also apply to fungi. However, a fungal *de novo*-maintenance pair has not been functionally or biochemically identified. Our studies identified one-half of this pair, Dnmt5, in the human pathogenic yeast *C. neoformans*. To our surprise, Dnmt5 is a maintenance enzyme *in vitro* with extraordinarily high specificity for hemimethylated substrates. Dnmt5 is not closely related to the conserved Dnmt1 maintenance methylase, and unlike Dnmt1, Dnmt5 requires ATP for activity and displays considerably higher specificity for hemimethylated DNA than Dnmt1 (Bashtrykov et al., 2014). Nonetheless, the chromatin signals that promote maintenance methylation in *C. neoformans* are shared with mammals and plants. Dnmt5 activity is strongly promoted by H3K9me, which it recognizes directly via its chromodomain and indirectly via an interaction with the Cryptococcus Swi6/HP1 which we show is a H3K9me reader as expected. Moreover, like Dnmt1, the activity of Dnmt5 is promoted by a Uhrf1-like protein that selectively recognizes hemimethylated DNA.

### Ancient loss of a *de novo* enzyme yet persistence of 5mC

Given that Dnmt5 is a highly-specific maintenance enzyme *in vitro*, it is unexpected that Dnmt5 is the only predicted cytosine methylase encoded by the *C. neoformans* genome. One possibility would be that Dnmt5 acts differently *in vivo*. However, we observed that methylation loss after depleting cells of Dnmt5 was not restored upon re-introduction of the enzyme. This was true even if Dnmt5 was introduced through a genetic cross. Tandem repeats are silenced in *C. neoformans* during sexual reproduction in a process called Sex-Induced Silencing (SIS) (Wang et al., 2010). However, SIS is an unstable RNAi-based phenomenon that does not result in methylation of repeats (Wang et al., 2010). As there does not appear to be a MIP-like phenomenon occurring in *C. neoformans*, it seems highly unlikely that Dnmt5 specificity is changed dramatically premeiotically to enable efficient *de novo* methylation of repeats. Instead, it appears that there is no efficient *de novo* methylation mechanism in this organism. Consistent with these observations, we determined that the ancestor of the pathogenic *Cryptococcus* species complex harbored a second DNMT homolog, DnmtX, whose gene was lost 150-50MYA. Our functional experiments demonstrate that extant DnmtX from three different species are each capable of triggering widespread DNA methylation *de novo* in *C. neoformans*, indicating that DnmtX is a *de novo* enzyme. Thus, we conclude that 5mC has endured in *C. neoformans* for millions of years through the action of Dnmt5.

### Inheritance, variation, and selection in an epigenome

Although we did not observed restoration of 5mC globally when Dnmt5 was removed and re-introduced, we did observe two apparent *de novo* events in two independent experiments. A caveat of this experiment is that cells that lack methylation to begin with would not be able to recruit Uhrf1 efficiently and thus may not well represent wild-type. Thus, we performed an experimental evolution experiment in wild-type cells. This work revealed preservation of 5mC patterns over an experiment of more than 120 generations, suggesting a high fidelity of inheritance for an epigenetic mark. Loss events occurred and considerably rarer gain events were also observed. The rare gain events may be produced by the action of Dnmt5 on unmethylated DNA, the activity of some other cellular methylase or gene conversion events that produce hemimethylated heteroduplex DNA that would then be a favored substrate for Dnmt5. The latter is plausible as the nonreciprocal transfer of 5mC has been observed during meiosis in *A. immersus* (Colot et al., 1996). Regardless of their source, these gain events, which seem to be rare and randomly distributed, provide a mechanism to inject new sites of methylation into the lineage. Because loss events are more frequent than gain events, we infer that the system is not at equilibrium during laboratory evolution, implying that natural selection is required to maintain long-term steady state level of 5mC. Further evidence for natural selection comes from our long read-assembled genomes and epigenomes of eight isolates of *C. neoformans var. grubii* that span the four major clades of this pathogen. These studies demonstrated higher levels of 5mC in centromeric transposon sequences than in non-transposon centromeric sequences. These data are compatible with a model in which 5mC loss and gain events in *C. neoformans* act in a manner akin to mutation in DNA sequence by producing random variation on which natural selection for silencing transposons acts.

In principle, if selection for 5mC at sites were sufficiently strong and if the fidelity of its replication were sufficiently high, 5mC patterns could be propagated unchanged for long periods of time. It is notable that we observe evolutionary conservation of 5mC position in our two-way analysis and that there are CG sites in Tcn3 elements that are shared between isolates that have diverged by ∼5 MY. Given the data at hand, a speculative possibility is that this sharing reflects an epigenetic equivalent of “identity-by-descent” in which methylation at specific sites has been maintained since the divergence of the compared strains purely through inheritance and selection (Thompson, 2013). The development of methods to generate an accurate phylogeny of transposon epigenomes and/or to align repetitive centromeric sequences might enable a determination of whether conservation of 5mC sites we observe is produced by such a mechanism. More generally, such methods might enable us to estimate the potential timescales of 5mC inheritance (epigenetic memory). We have not observed any sequence specificity for Dnmt5 based on searches for motifs, and we found introduced *in vitro* methylated DNA to be well maintained by Dnmt5 *in vivo*. This argues that conservation of methylation sites we observe is not explained by a detectable requirement by Dnmt5 for particular sequences beyond hemimethylated CG dinucleotides.

### Evolution of DNA methylation: suppressors and drivers of loss

Our phylogenetic analysis shows that that human pathogens *C. neoformans var. grubii*, *C. neoformans var. neoformans*, and the *C. gattii* descended from an ancestor that lost DnmtX. It has been suggested that two mechanisms enable gene loss during evolution: mutational robustness (produced by redundant pathways) and environmental variability (Albalat and Canestro, 2016). While these concepts may be germane to understanding how the loss of DnmtX was tolerated, the fact that 5mC was retained after the loss raises a third possibility, namely a long-lived phenotypic lag. That is, if methylation is effectively maintained by Dnmt5 after loss of DnmtX, this might enable suppressors to evolve to compensate the loss. Such suppressors may involve changes in transposon content that in some way eliminate the need for a dedicated *de novo* system or changes in RNAi mechanisms that we and others have shown suppress transposon movement in *C. neoformans* (Burke et al., 2019; Janbon et al., 2010). Conversely, it is possible that there could be benefits to losing DnmtX, particularly if it were to be involved in a MIP-like phenomenon, which would limit evolution by gene duplication and divergence. The loss of a half a *de novo*-maintenance system pair raises the possibility that this represents an intermediate step that enables (via a phenotypic lag) the loss of cytosine methylation in a lineage, a common evolutionary occurrence.

## EXPERIMENTAL PROCEDURES

### Yeast growth and manipulations

Yeast manipulation and protocols were performed as described in (Chun and Madhani, 2010). Tagging, deletions and insertions of genes were obtained through homologous recombination by transforming about 10 µg of either a PCR product or a PmeI-digested plasmids bearing the sequence to be transformed. The *RI-DMT5* strain used for WGBS was produced by deleting the 5’ portion of the *DMT5* gene and then restoring it by homologous replacement. To be certain that no residual DNA methylation was present in *dmt5Δ* prior to introduction of the targeting construct that restores the gene, the *dmt5Δ* strain was first propagated in YPAD for at least 90 generations.

To express DnmtX in an *RI-DMT5* strain, genomic DNA from *K. pini, K. mangroviensis* and *K. bestiolae* was used as a template to amplify the genes putative DnmtXs (*K. mangroviensis* I_20_3_05465, *K. pini* I302_00877, *K. bestiolae* I_20_6_02865). DnmtX was cloned downstream a *GAL7* promoter and the entire construct was then inserted at the *URA5* locus. Upon transformation, cells were selected in YPAD+nourseothrycin and then propagated for two sequential single-colony streaks in presence of galactose to induce DnmtX.

Other strains generated by transformation were colony-purified, patched, verified, and frozen. Liquid cultures were inoculated directly from the frozen stock. A list of strains used in this study can be found on Table S2.

### Genetic crosses

Crosses were carried out using strains of the KN99 background. Cells of different mating types were grown overnight in YPAD, spotted on mating plates (4% Agar, 10% 10X Murashige and Skoog basal medium, 0.1mg/ml myo-inositol) and kept in the dark for 1 to 2 weeks. Spores were collected with a toothpick and resuspended in 500 µl of water. Dilutions of this suspension were then spread on selective media.

### DNA extraction and Southern analysis

DNA from *C. neoformans, C. pinus, K. mangroviensis* and *C. bestiolae* was extracted as previously described (Chun and Madhani, 2010) with some modifications. In brief, 100 OD of yeast (100 ml, OD1) were harvested, frozen in liquid nitrogen, resuspended in 5ml CTAB buffer (100 mM Tris pH 7.6, 1M NaCl, 10 mM EDTA, 1% Cetyltrimethyl ammonium bromide, 1% ß-mercaptoethanol) and incubated at 65°C for at least 1h. 3ml of chloroform were added to the lysate, mixed and spun for 5min at 3000g. The aqueous phase was then precipitated by adding the same volume of isopropanol. The dried pellet was resuspended in 300 µl TE containing 1 µg RNAse A (Thermo Scientific) and incubated 30 min at 37°C followed by addition of 3 µl Proteinase K (20 mg/ml) and incubation at 50°C for 1 h. Phenol-chloroform and chloroform extraction was carried out and the DNA is finally precipitated by addition of 1/10 volume NaOAc and three volumes of ethanol. The DNA was additionally purified with Genomic DNA Clean & Concentrator (D4065, Zymo Research). For Southern analysis, 10µg were then digested with the meCpG sensitive enzyme HpyCH4IV (NEB), separated by electrophoresis on a 1% agarose gel and transferred to nylon membrane (Hybond NX, Amersham) by capillary action. DNA was crosslinked to the membrane using UV light. PCR products were used as template to incorporate radioactive ^32^P-dCTP (Perkin Elmer) using the High Prime kit (Roche) according to manufacturer’s protocol. Primer sequences for the probes are listed in Table S3.

### Stress conditions experimental procedures

To assess if specific stress conditions induce *de novo* DNA methylation activity, wild-type and RI-Dnmt5 cells were grown in the presence of stressors. 2.3 M NaCl at 30°C for 3h, 140 mM/ 2.3mM NaNO2/succinic acid at 30°C for 3h, 0.14% H_2_O_2_ at 30°C for 3 h, 0.9% SDS at 30°C for 3h, 1.9 mg/ml caffeine at 37°C for 20h, DMEM at 37°C, 5% CO_2_, for 24 h. The concentrations of stressors used represent the maximum concentration that allows growth in the conditions tested.

### Protein extraction and western analysis

Two millilitres of culture at OD_600_=1 were collected by centrifugation and frozen in liquid nitrogen, resuspended in 10% TCA and incubated on ice for 10min. Cells were washed once with acetone and air-dried for 10min. The pellet was then resuspended in 150 µl 2x Laemmli buffer adjusted with 80 µl Tris-HCl (pH 8.0) and bead-beated 2 x 90 s. The lysate was boiled for 5min and centrifuged to remove residual cells. Western analysis was performed using anti-FLAG antibody (1:3000-F3165, sigma), anti-HA (1:10000-26183, Thermo Fisher) or anti-H3 antibody (1:1000-PA5-16183, Thermo Fisher) diluted with 5% milk in TBS-T (10 mM Tris-HCl pH 7.6, 150 mM NaCl, 0.1% Tween 20) for 1h followed by 2 x 10 min washes in TBS-T. The membrane was then incubated with antibody anti-rabbit or anti-mouse conjugated to HRP (BIO-RAD) (1:8000 in 5% milk+TBS-T) for 45min followed by 2 x 10 min washes in TBS-T. The membrane was then incubated in SuperSignal West pico luminol (Thermo Scientific) and visualized using film.

### Affinity purification and mass spectrometry analysis

Dynabeads M-270 Epoxy (Cat n. 14301) were conjugated following the manual with the following modifications. 150 μg anti-FLAG antibody (Sigma-F3165) was conjugated with 1×10^9^ beads overnight at 37°C. The beads were washed once with 100 mM Glycine HCL pH 2.5, once with 10mM Tris pH 8.8, 4x with 1x PBS for 5 min, once with PBS + 0.5% Triton X-100 for 5 minutes, once PBS +0.5% Triton X-100 for 15 min. Beads were the resuspended in the PBS and kept at 4°C before using them.

4 L of *C. neoformans* cells were grown on YPAD at 30°C until OD2, harvested, resuspended in 10 ml Lysis buffer (25 mM HEPES-KOH, pH 7.9, 2 mM MgCl_2_, 100 mM KCl, 16% glycerol, 0.1% Tween 20) and dripped into liquid nitrogen to form ‘popcorn.’ Cell were lysed by cryo-grinding in a SPEX Sample Prep 6870 instrument for 3 cycles of 2 min at 10 cps with 2 min of cooling between each cycle. The extract was then thawed at 4°C, the final volume was brought to 80 ml with Lysis Buffer+2 mM CaCl_2_ and incubated with 1000U of RQ1 DNase (Promega-PRM6101) for 30min at RT. At this point, the extract was clarified by centrifugation at 40Kxg for 40 min at 4°C and FLAG-conjugated magnetic beads were added to the supernatant and incubated for 4h at 4°C with rotation. The beads were then washed 3×10min with lysis buffer and using a magnetic rack. Bound proteins were eluted three times in 150 µl elution buffer (25 mM HEPES-KOH, pH 7.9, 2 mM MgCl_2_, 300 mM KCl, 20% glycerol) with 1 mg/ml 3X FLAG peptide (Sigma). For the co-immunoprecipitation experiments, 1 L culture was used in the assay and the final elution step was performed by boiling the beads with 2x Laemmli buffer for 10 min.

Label free mass spectrometry was performed as follows. Samples were precipitated by methanol/ chloroform. Dried pellets were dissolved in 8 M urea/100 mM TEAB, pH 8.5. Proteins were reduced with 5 mM tris(2-carboxyethyl)phosphine hydrochloride (TCEP, Sigma-Aldrich) and alkylated with 10 mM chloroacetamide (Sigma-Aldrich). Proteins were digested overnight at 37 °C in 2 M urea/100 mM TEAB, pH 8.5, with trypsin (Promega). Digestion was quenched with formic acid, 5 % final concentration.

The digested samples were analyzed on a Fusion Orbitrap tribrid mass spectrometer (Thermo). The digest was injected directly onto a 25 cm, 100 □m ID column packed with BEH 1.7□m C18 resin (Waters). Samples were separated at a flow rate of 300 nl/min on a nLC 1000 (Thermo). Buffer A and B were 0.1% formic acid in water and 0.1% formic acid in acetonitrile, respectively. A gradient of 1-25% B over 110 min, an increase to 40% B over 10 min, an increase to 90% B over 10 mins and held at 90%B for a final 10 min was used for 140 min total run time. Column was re-equilibrated with 20 □l of buffer A prior to the injection of sample. Peptides were eluted directly from the tip of the column and nanosprayed directly into the mass spectrometer by application of 2.5 kV voltage at the back of the column. The Orbitrap Fusion was operated in a data dependent mode. Full MS scans were collected in the Orbitrap at 120K resolution with a mass range of 400 to 1500 m/z and an AGC target of 4e^5^. The cycle time was set to 3 sec, and within this 3 sec the most abundant ions per scan were selected for CID MS/MS in the ion trap with an AGC target of 5e^4^ and minimum intensity of 5000. Maximum fill times were set to 50 ms and 100 ms for MS and MS/MS scans respectively. Quadrupole isolation at 1.6 m/z was used, monoisotopic precursor selection was enabled and dynamic exclusion was used with exclusion duration of 5 sec.

Protein and peptide identification were done with Integrated Proteomics Pipeline – IP2 (Integrated Proteomics Applications). Tandem mass spectra were extracted from raw files using RawConverter (He et al., 2015) and searched with ProLuCID (Xu et al., 2015)against *Cryptococcus neoformans* UniProt database. The search space included all fully-tryptic and half-tryptic peptide candidates. Carbamidomethylation on cysteine was considered as a static modification. Data was searched with 50 ppm precursor ion tolerance and 600 ppm fragment ion tolerance. Identified proteins were filtered using DTASelect (Tabb et al., 2002) and utilizing a target-decoy database search strategy to control the false discovery rate to 1% at the protein level (Peng et al., 2003).

Proteins that did not appear in the untagged control that displayed at least 10% sequence coverage in both duplicates were considered hits in our analysis. Common contaminants (e.g. ribosomal proteins) were removed from the analysis. The complete dataset can be found in Table S4.

### Bisulfite conversion and library construction

Bisulfite library construction was performed as previously described (Urich et al., 2015). Briefly, 100 OD units of yeast were harvested, lyophilised overnight and the DNA was extracted using DNeasy Plant Mini Kit (Qiagen). The DNA was additionally purified with a Genomic DNA Clean & Concentrator (D4065-Zymo Research). Genomic DNA was quantified with Qubit dsDNA BR Assay Kit (ThermoFisher). 5ng of lambda DNA (Promega) was added to ∼1ug yeast DNA and fragmented using a Bioruptor Pico instrument (Diagenode-13 cycles, 30s/30s On/Off). The DNA was end-repaired and ligated to methylated adapters. Two rounds of bisulfite conversion of the adapter-ligated DNA were performed using EZ-DNA methylation Gold Kit (D5005-Zymo Research), and used as template for three separate PCR reactions (eight cycles of amplification) as described in (Urich et al., 2015). DNA obtained from the three PCR reactions were then combined and analyzed for size on a High Sensitivity DNA chip (Agilent Technologies) and sequenced as SE100. Reads from the bisulfite-treated library were trimmed using cutadapt (https://cutadapt.readthedocs.io/en/stable/) and analyzed with Bismark (Krueger and Andrews, 2011) using bowtie2 with the options: --score_min L,0,0. Unconverted reads were removed using Bismark-filter_non_conversion and methylation was called using the standard setting.

To quantify CG methylation levels globally, data were further filtered using a custom Python script for signals at CG dinucleotide that displayed a fractional methylation of 0.5 or greater on both strands.

### Chromatin immunoprecipitation

ChIP and library preparation were performed as previously described (Homer et al., 2016) with the following modifications. Cells were lysed using a bead beater (8 cycles x 90 s) and lysate was clarified by centrifugation 10 min at 6800 g. The chromatin pellet was sonicated in a Bioruptor Pico instrument (Diagenode; 25 cycles, 30s/30s on/off) and centrifuged for 20 min at 20000 g. The cleared lysate was incubated overnight with 6 µg anti-histone H3K9me2 (ab1220, Abcam) or 4 µl anti-FLAG antibodies (F3165, sigma) with the addition of 30 µl of protein A (for anti-H3K9me) or protein G (for anti-FLAG) Dynabeads (Thermo Fisher). Library preparation was performed as described in (Homer et al., 2016). Reads were trimmed for adaptor sequence (“GATCGGAAGA”) and filtered for reads of at least 50 nt in length using Cutadapt (Martin, 2011) with were aligned to the C. neoformans genome using bowtie1 (modified parameters: -v2, -M1, --best) (Langmead et al., 2009). Alignment files were processed with SAMtools (Li et al., 2009a) and converted to bedgraph format for visualization with BEDTools (Quinlan and Hall, 2010). Bedgraphs were scaled by million aligned reads and normalized to whole cell extract at each genomic position, then smoothed using a 500 bp centered rolling mean using custom Python scripts.

For scatter plots, reads (were counted in each region (centromere or telomere) using the Pysam Python library (https://github.com/pysam-developers/pysam). RPKM was determined by scaling the reads in each region, *n,* to the total number of aligned reads in the sample, *M* (in millions), and the length of the region, *l* (in kb):

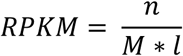

Normalized enrichment in ChIP over whole cell extract, *E*, was calculated based on RPKM as follows:

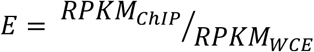

### Methylated DNA immunoprecipitation (MeDIP)

The initial steps are the same as described for the bisulfite library protocol (Urich et al., 2015) with the exception that for MeDIP, unmethylated adapters were ligated to repaired, dA-tailed DNA. Upon ligation and purification, DNA was boiled for 10 min and kept on ice for 5min. Ice cold binding buffer (10 mM sodium phosphate buffer, pH 7, 140 mM NaCl 0.05% Triton X-100) was added to the DNA together with 1 µl of anti-5-methylcytosine antibody (A-1014-050, Epigentek) and incubated on a nutator overnight at 4°C. 20 µl Protein G Dynabeads (Thermo Fisher) were added and incubated on a nutator for further 2 h at 4°C. Beads were washed 10×5 min in binding buffer and the DNA was eluted by incubating the beads with Elution Buffer (TE, 0.25 mg/ml Proteinase K, 0.25% SDS) for 2h at 55°C. The DNA was purified with NucleoSpin PCR clean-up columns (Macherey-Nagel) using NTB buffer (Macherey-Nagel) and PCR amplified as for ChIP. Bioinformatics analyses were performed as for ChIP-seq.

### *In vitro* methylation and 5mC-qPCR assay

The construct was *in vitro* methylated with HpaII methyltransferase (NEB), which recognises the CCGG sequence and modifies the internal cytosine residue. At least 10 µg of the targeting construct were incubated with 20U HpaII methyltransferase for 2 h at 37°C followed by inactivation of the enzyme at 65°C for 20 min. The DNA was then purified with NucleoSpin Gel and PCR Clean-up Columns (Macherey-Nachel, 740609) and the efficiency of the *in vitro* methylation reaction was assayed by digesting 100 ng of the in vitro methylated product with 1U of the HpaII endonuclease (run: undigested, unmethylated+digestion, methylated+digestion, Figure 5C). Transformation was carried out with the *in vitro* methylated or with the unmodified fragment and the transformants were selected in the presence of G418.

To assess the ability of cells to maintain 5mC at the *in vitro* methylated CCGG sites, we calculated the methylation fraction for specific sites as ratio of qPCR values across the CCGG-site of interest (5mC protects from restriction enzyme digestion) to a region that does not contain the CCGG site. To test the capability of Dnmt5 to promote the spread 5mC, adjacent sites to HpaII sequence were used and in specific CGCG sites (BstUI) or CGWCG sites (Hyp99I). Genomic DNA was prepared as previously described and cleaned up with a Genomic DNA Clean & Concentrator kit (D4065-Zymo Research). 50 ng of genomic DNA was digested with the appropriate restriction endonuclease and purified using Genomic DNA Clean & Concentrator kit (D4065-Zymo Research). 0.6 ng was used as template for qPCR analysis using PowerUp SYBR Green Master Mix (Life Technologies) with primers listed in Table S3.

### Experimental evolution and 5mC analysis

Wild-type cells from a starter culture (g0) were propagated at 30°C in rich media for 120 generations maintaining the logarithmic growth by diluting the culture as soon as it reached an OD600 of 1. After 120 generations, the cells were plated on YPAD agar plates to form single colonies at 30°C. Two independent isolates (colony 1 and colony 2) were grown overnight in rich media to OD600 of 1 and the genomic DNA extracted processed in technical duplicates for WGBS. WGBS libraries and analysis were performed as described above.

Differential methylation site (DMS) analysis was performed using CGmap tools (https://cgmaptools.github.io). Bismark outputs were converted into CGmap file using cgmaptools-convert-bismark2cgmap and then sorted and intersected with cgmaptools-sort and – intersect, respectively. Differential analysis was performed using cgmaptools-dms on the intersected files. Only sites of symmetric CG with >50% methylation that were reproducible between technical replicates were considered in the analysis. DMS was considered significant when p-value<0.05.

### Recombinant protein expression and purification

A codon-optimized DNA sequence encoding *C. neoformans* Dnmt5 (residues 1-150) was cloned into pETARA or pMAL vectors and used to transform *E. coli* strain BL21(DE3) (Harris et al., 2001; Reményi et al., 2005). Transformed cells were grown to OD_600_ = 0.8 in 2x YT medium, then induced with 1 mM IPTG overnight at 18°C. Recombinant GST-Dnmt5(1-150)-6xHis was purified with Ni-NTA agarose resin (Qiagen), measured by A_280_ (ε = 66,030 cm^-1^ M^-1^), and used for histone peptide array binding assays. Recombinant MBP-Dnmt5(1-150)-6xHis was purified with Ni-NTA agarose resin (Qiagen), measured by A_280_ (ε = 89,590 cm^-1^ M^-1^), and used for fluorescence polarization binding assays.

A codon-optimized DNA sequence encoding full-length *C. neoformans* Uhrf1 was cloned into pBH4 and expressed in *E. coli* as above. Recombinant 6xHis-Uhrf1 was purified with Ni-NTA agarose resin (Qiagen), measured by A_280_ (ε = 40,920 cm^-1^ M^-1^), and used for electrophoretic mobility shift assays.

Full-length cDNA encoding Swi6 was cloned from *C. neoformans* and expressed in *E. coli* as above using the pBH4 vector. Recombinant 6xHis-Swi6 was purified with Ni-NTA agarose resin (Qiagen), measured by A_280_ (ε = 45,330 cm^-1^ M^-1^), and used for fluorescence polarization binding assays.

For expression in *S. cerevisiae*, full-length cDNA encoding Dnmt5 was cloned from *C. neoformans*, inserted into the 83ν vector(Li et al., 2009b), and used to transform the *S. cerevisiae* strain JEL1(Lindsley and Wang, 1993). Starter cultures were grown overnight in SC -ura medium (2% glucose), then used to inoculate 2 L cultures of YPGL medium (1x YEP, 1.7% lactic acid, 3% glycerol, 0.12% glucose, 0.15 mM adenine) to a starting OD_600_ of 0.03. After growth at 30°C to an OD_600_ of 1.0, expression was induced by addition of 2% galactose. After 6 hr of continued growth at 30°C, cells were harvested, washed once in TBS (50 mM Tris-Cl pH 7.6, 150 mM NaCl), and snap frozen. Frozen cells were lysed in a ball mill (6x 3 min at 15 Hz), resuspended in Ni-NTA lysis buffer (50 mM NaH_2_PO_4_ pH 8, 300 mM NaCl, 10% glycerol, 10 mM imidazole, 2 mM β-mercaptoethanol, 0.02% NP-40, 1x CPI), and centrifuged 20,000 x *g* for 30 min at 4°C. Lysate was bound to Ni-NTA resin in batch format for 2 hr at 4°C, after which resin was washed in column format with five bed volumes Ni-NTA buffer followed by ten bed volumes Ni-NTA wash buffer (identical to Ni-NTA lysis buffer except 20 mM imidazole). Finally, bound protein was eluted with four bed volumes of elution buffer (identical to Ni-NTA lysis buffer except 300 mM imidazole and no detergent). Protein was dialyzed against storage buffer and applied to a HiTrap Q HP anion exchange column (GE Life sciences) pre-equilibrated in buffer (50 mM HEPES-KOH pH 7.9, 150 mM KCl, 10% glycerol, 2 mM β-mercaptoethanol). Fractions were collected across a 150-1000 mM KCl gradient, and those containing Dnmt5 were pooled, dialyzed against storage buffer, and frozen.

### Histone peptide array binding assay

Modified histone peptide arrays (Active Motif) were blocked with 5% dried milk in TBS-T (10 mM Tris-HCl pH 7.6, 150 mM NaCl, 0.1% Tween 20) overnight at 4°C. Each slide was subsequently washed in TBS-T (4×5 min) and binding buffer (2×5min). Binding buffer consisted of 50 mM HEPES-KOH pH 7.9, 200 mM NaCl. The slide was incubated with 0.5 μM GST-Dnmt5(1-150) in binding buffer for 4 hr at 23°C. It was then washed with TBS-T (4x 5 min). Next, the slide was incubated with anti-GST antibody (Sigma G7781; 1:13,000 dilution) in 5% dried milk in TBS-T for 1hr at 23°C. After washing in TBS-T (4x 5 min), the slide was incubated with HRP-conjugated goat anti-rabbit secondary antibody (sc-2004, Santa Cruz Biotechnology; 1:5,000 dilution) in 5% milk in TBS-T for 1hr at 23°C. The slide was washed in TBS-T (4x 5 min) and visualized using SuperSignal West Pico chemiluminescent substrate (Pierce) and a ChemiDoc MP System (BioRad). Signal intensity normalization was performed using Array Analyze Software (Active Motif).

### Fluorescence polarization binding assay

Peptides were synthesized by GenScript (Piscataway, NJ) or Peptide 2.0 (Chantilly, VA) to >95% purity. For study of H3K9 methylation, unlabeled peptides corresponded to residues 1-15 of *C. neoformans* histone H3. For study of H3K27 methylation, unlabeled peptides corresponded to residues 23-34 of *C. neoformans* histone H3 followed by a cysteine residue. Peptide concentrations were determined by A_205_ or, in the case of fluorescein-conjugated peptides, A_495_ (ε = 80,000 M^-1^ cm^-1^).

For direct measurement of peptide binding, increasing concentrations of MBP-Dnmt5(1-150) were incubated with 10 nM labeled peptide in a solution of 20 mM HEPES pH 7.9, 120 mM KCl, 0.8 mM DTT, and 0.01% NP-40. Peptide fluorescence polarization was measured using a Spectramax M5e plate reader (Molecular Devices) and non-stick 384-well plates (Corning 3820). For competition assays, unlabeled competitor peptides were added at increasing concentration in the setting of 5 μM MBP-Dnmt5(1-150) or 30 μM 6xHis-Swi6 and 10 nM fluorescein-conjugated H3K9me3 peptide. Dissociation constants were calculated using a competition binding equation in Prism (GraphPad Software)(Pack et al., 2016):

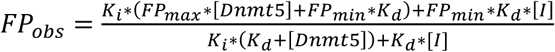

Dissociation constants for H3K9me3 peptide were comparable when measured directly (1.3 μM) or as a competitor (1.5 μM).

### Gel mobility shift assay

Primer sequences are listed in Table S3 and were annealed to generate unmethylated, hemimethylated, or symmetrically methylated 20 bp dsDNA substrates. The substrates were then radiolabeled using the KinaseMax kit (Ambion) and ATPγ-^32^P (Perkin Elmer), after which they were purified using a G-25 illustra microspin column (GE Life Sciences). For direct measurement of DNA binding, recombinant 6xHis-Uhrf1 was incubated with 0.2 nM labeled DNA probe in a 10 μl solution of 15 mM HEPES-KOH pH 7.9, 7.5% glycerol, 75 mM KCl, 0.075% NP-40, 0.05 μg/ul ^p^oly-dI:dC (Sigma), 0.5 μg/ul BSA, 1 mM DTT, and 5 mM MgCl_2_. After 30 min equilibration at 23°C, samples were resolved in polyacrylamide gels (4.5% acrylamide:bis 29:1 (Bio-Rad), 1% glycerol, 0.25X TBE) at 4°C. Gels were subsequently dried and imaged using a storage phosphor screen (Amersham) and a Typhoon 9400 imager (Amersham). Densitometry was performed using ImageJ. For competitive binding assays, conditions were as above except Uhrf1 protein was kept constant at 150 nM in the presence of 0.2 nM labeled hemimethylated probe and excess amounts of unlabeled DNA oligonucleotides.

### DNA methyltransferase assay

DNA oligonucleotides were synthesized and annealed to generate 20 or 60 bp dsDNA substrates (Table S3). DNA methylation was monitored in multiple turnover conditions by incubating 30 nM Dnmt5 in a solution of 50 mM Tris pH 8, 25 mM NaCl, 10% glycerol, and 2 mM DTT, in the ^pr^esence of 5 μM DNA substrate. When indicated, ATP and MgCl_2_ were supplemented at 1 mM, histone tail peptides corresponding to histone H3 residues 1-15 were added at 5 μM, Uhrf1 was added at 4 μM, and Swi6 was added at 4 μM. Reactions were initiated by the addition of 4 μM ^3^H-SAM (Perkin Elmer) and carried out at 23°C. Reaction aliquots were removed at indicated time points and quenched in a solution of 10 mM SAM in 10 mM H_2_SO_4_. The quenched solution was pipetted onto DE81 filter paper and allowed to dry for 10 min. Filter papers were subsequently washed three times in 200 mM ammonium bicarbonate (5 min each), then once with water (5 min). Filter papers were then rinsed twice in ethanol, after which they were dried for 20 min. Filters were added to scintillation fluid (Bio-Safe NA, Research Products International Corp.), and bound ^3^H was detected in an LS 6500 scintillation counter (Perkin Elmer). Background signal was assessed using reactions that lacked Dnmt5 enzyme. Detectable signal was defined as reactions that exhibited cpm greater than 2-fold above background at each measured time point. Background signal was typically 50-100 cpm and signal for productive reactions ranged from ∼1000 to ∼100,000 cpm depending on conditions and time point. For productive reactions, separate experiments were performed to confirm that the DNA substrate was at saturating concentration. For reactions in which methylation was detected, rates were calculated over the first 15-20 min where reaction progress was linear and less than 10% of available hemimethylated sites had been acted upon. These initial rate values were divided by Dnmt5 concentration to obtain k_obs_. DNA substrate was confirmed to be present at saturating concentration for these experiments. Serial dilutions confirmed that ^3^H detection using DE81 was linear to the level of background signal.

### Phylogenetic methods

For phylogenetic analysis, orthologs were identified across sequenced Tremellales (Cuomo, manuscript in prep) based on BLASTP pairwise matches with expect< 1e-5 using ORTHOMCL v1.4.(Li et al., 2003) A phylogeny was estimated from 4080 single copy genes as follows. Individual proteins were aligned using MUSCLE (Edgar, 2004) and the individual alignments were concatenated and poorly aligning regions removed with trimal.(Capella-Gutierrez et al., 2009) This sequence was input to RAxML (Stamatakis, 2006) version 7.7.8 (raxmlHPC-PTHREADS-SSE3) and a phylogeny estimated in rapid bootstrapping mode with model PROTCATWAG and 1,000 bootstrap replicates.

### Nanopore sequencing and bisulfite analysis of eight *C. neoformans var grubii* isolates

Genomic DNA was extracted as described above with the only modification that the cells were incubated with lysis buffer for only 30 min at 65°C and other 30 min at room temperature.

Oxford Nanopore libraries were constructed using the 1D ligation kit (SQK-LSK108) and loaded on a FLO-MIN106 flow cell for each sample on a Minion. Basecalling was performed using Albacore v2.0.2. A total of 63- to 314-fold average genome coverage was generated for each of the eight samples. ONT fastq reads were assembled using Canu(Koren et al., 2017) release v1.5 with the following parameters: -nanopore-raw <input.fastq>, correctedErrorRate=0.075, and stopOnReadQuality=false. The Canu1 assembly was corrected using nanopolish (v0.8.1); reads were aligned to the Canu assembly using bwa-mem2(Li, 2013) (0.7.12) with the “-x ont2d” parameter and nanopolish (https://github.com/jts/nanopolish) v0.9 variants in --consensus mode was run on each contig to generate a polished consensus. Next, assemblies were manually inspected and edited as follows. Contig ends without telomere repeats were aligned using NUCmer3(Kurtz et al., 2004) to unassembled contigs and extended where alignments were uniquely mapped and contained both contig ends. Redundant small contigs with high identity NUCmer alignments to larger contigs were examined for read support; while all appeared redundant, some of these regions had higher accuracy and were used to correct the large contigs in two assemblies. Two manual joins between contigs were also made where supported by alignments to unassembled contigs. Alignment to the H99 genome4(Janbon et al., 2014) with NUCmer was also used to validate these assemblies. All 8 assemblies showed the same rearrangement between H99 chromosomes 3 and 11; while seven of these assemblies appeared co-linear, Bt106 showed a unique rearrangement between H99 chromosomes 4 and 10 that was validated by read alignments. Finally, Illumina paired reads were aligned to the resulting polished contigs using bwa mem (version 0.7.7-r441) followed by Pilon5(Walker et al., 2014) (version 1.13) correction using the --fix all setting; this step was repeated for a second round of alignment and Pilon polishing.

### Data access

All raw Oxford nanopore sequence and genome assemblies are available in NCBI under BioProject PRJNA517966.

WGBS library preparation was performed as described above and sequenced as SE150. Two technical replicates of the same genomic DNA were prepared for each sample. Reads from the bisulfite-treated library were analyzed in parallel with Bismark (Krueger and Andrews, 2011) and BSseeker2(Guo et al., 2013). Bismark alignment: bowtie2 with the options: --score_min L,0,0. Unconverted reads were removed using Bismark-filter_non_conversion and methylation was called using the standard setting. BSseeker2 alignment: bowtie2, --mismatches=0 --XS=0.5, 5, call methylation: -x.

Bismark outputs were converted into CGmap file using cgmaptools-convert-bismark2cgmap (https://cgmaptools.github.io). The CGmap files from the technical replicates from BSseeker2 and Bismark were then cross-compared and only symmetric methylated CGs with at least 50% methylation and covered by >=2 reads were considered in further analysis.

5mC calls on the Nanopore long reads data was performed using Nanopolish(Simpson et al., 2017) call-methylation and scripts/calculate_methylation_frequency.py.

The final table obtained was then converted into a CGmap file to be compared with our WGBS analysis. Since the calling 5mC with Nanopolish cannot distinguish methylated status of those CGs within 10bp of each other, the same methylation status was assigned to the entire group in the final CGmap. This results in a modest overestimation of the overall methylation status.

### Analysis of 5mC conservation

Centromeres were pairwise-aligned to a reference (A1-35-8) using NUCmer and only the longest increasing subset was kept for each alignment. Methylation calls from WGBS (for whole genome methylation analysis) or WGBS-combined with ONT-methylation calls (for methylation analysis on transposons) were then projected onto these coordinates.

For the whole genome methylation analysis (Figure 7D), we binned sites with conserved CG sequence in X number of strains (from 2 to 8) into CG conservation groups and asked the relative rates of methylation in each bin. A null distribution was derived by permuting within the CG conservation group to mimic the heterogeneity in methylation expected given sequence conservation. For example, to generate the null distribution for strain Bt106 where 4 sites are shared, all the CGs that Bt106 shares with 3 other strains were permuted. This resulted in a much more conservative estimate than the fully random model.

For the Tcn analysis, first a class was of transposable elements was chosen (e.g. Tcn3), then the longest full-length Tcn in that class was chosen as a reference. Each Tcn was then aligned to the reference using Bioconductor pairwiseAlignment in global mode. WGBS were used to evaluate the methylation status. In cases where WGBS was not available (e.g. repetitive regions), we used the ONT calls to fill in the missing information. This is justified by high concordance between the two methods where there is bisulfite data (calculated as Pearson coefficient - Figure S12E).

Independence of methylation and sequence was tested by computing a goodness-of-fit at each site relative to the reference, conditioned on the aligned sequence. The procedure follows: 1) the expected value for each site was assumed to be the relative rate of methylation for the respective repeat (e.g. the random uniform model); 2) the statistic for each repeat at a specific C/G is (observed methylation status – repeat methylation rate) ^2/ repeat methylation rate; 3) the per repeat and site statistic was then the sum across all repeats and sites. The null distribution was generated by permuting methylation status within each repeat then recomputing the test statistic 1000 times.

### Liquid Chromatography Mass Spectrometry (LC-MS) method to quantify 5mC

Genomic DNA was isolated as previously described and purified with Genomic DNA Clean & Concentrator kit (D4065 - Zymo Research). Enzyme mix provided (New England Biolabs) was used to digest the gDNA to nucleosides. Standard nucleosides were diluted in different ratios using LC-MS water. Identical amount of isotope labelled-internal standards (provided by T. Carell) were spiked in the dilutions of standards and in the samples. Chromatographic separation of nucleosides was achieved using an Agilent RRHD Eclipse Plus C18 2.1 × 100 mm 1.8 μm column and the analysis was performed using an Agilent triple quadrupole mass spectrometer, (1290) UHPLC hyphenated to a (6490) QQQ-MS. The LC-MS method used for the quantitation of 5mC was performed as previously described (Leitch et al., 2013). A standard curve was generated by dividing unlabeled over isotope-labelled nucleosides and used to covert the peak area values to quantities. The signal-to-noise (S/N) used for quantification (above 10) was calculated using the peak-to-peak method. Global 5mC level is quoted relative to total levels of deoxyguanine (dG). The LOD (level of detection) for 5mC in the experiments was 25 amol and LOQ (limit of quantification) was 250 amol.

## Acknowledgements

ACKNOWLEDEMENTS

We thank all members of the members of the Madhani lab for scientific discussions and support, Yin Shen (UCSF) for methylated adaptors and advice on WGBS, and Matteo Pellegrini (UCLA) for advice on bioinformatics analysis of WGBS. We are thankful to John Perfect for the *C. neoformans var grubii* isolates. S.C. was supported by an EMBO Long term Postdoctoral Fellowship. Research in the Madhani laboratory is supported by grants from the U.S. National Institutes of Health. H.D.M. is a Chan-Zuckerberg Biohub Investigator. Mass spectrometry work in the Yates lab was supported by NIH grant 8 P41 GM103533. Work in Hajkova lab is supported by MRC funding (MC_US_A652_5PY70) and by an ERC grant (ERC-CoG-648879 – dynamicmodifications).

## SUPPLEMENTARY FIGURE LEGENDS

Figure S1. The chromodomain of Dnmt5 binds to H3K9me peptides

A. Coomassie Brilliant Blue staining of GST-Dnmt5(1-150) and MBP-Dnmt5(1-150) after affinity purification from lysates of *E. coli* and resolution by SDS-PAGE. This truncation contains the Dnmt5 chromodomain.
B. Dnmt5(1-150)-GST binding to a peptide microarray of human-derived histone sequences and modifications. Intensity represents background-subtracted signal at peptide array spots exhibiting the modifications indicated below the graph. Data are the average from two peptide arrays; error bars represent SE.
C. Binding of MBP-Dnmt5(1-150) to H3K27 peptides, as assessed by a fluorescence polarization binding competition assay. Dnmt5(1-150) was bound to a fluorescently-labeled H3K9me3 peptide in the presence of increasing concentrations of unlabeled H3K27me0/2/3 peptides corresponding to a region of *C. neoformans* H3. Polarization was normalized to that observed in the absence of competitor peptide; K_d_ calculated from 2-3 replicates.

Figure S2. Interactions of Dnmt5 and heterochromatin

A. ChIP-seq analysis of FLAG-Dnmt5 in wild-type and *clr4Δ* backgrounds compared to untagged strain. Shown are the data for chromosome 13.
B. Effect of chromodomain aromatic cage mutations on binding of MBP-Dnmt5(1-150) to H3K9 peptides. Wild-type or *W87AY90A* mutant MBP-Dnmt5(1-150) was incubated at increasing concentrations with H3K9me3 peptide. Polarization was normalized such that a value of 1 was equivalent to the polarization in the absence of MBP-Dnmt5(1-150) protein.
C. Coomassie staining of 6xHis-Swi6/HP1 after purification from *E. coli* and resolution by PAGE. Left: chromodomain and chromoshadow domains of full-length Swi6 are indicated. Right: binding of Swi6 to H3K9 peptides, as assessed by fluorescence polarization binding competition assay. Swi6 was bound to labeled H3K9me3 peptide in the presence of increasing concentrations of unlabeled H3K9me0/1/2/3 peptides. Polarization was normalized to that observed in the absence of competitor peptide; K_d_ represents average ± 95% CI of 2-3 replicates.
D. List of factors detected by Mass Spec analysis. The proteins used as baits are marked in yellow.
E. Western blot of co-immunoprecipitation assay performed using FLAG antibody that recognize FLAG-Dnmt5. The presence of Swi6-HA in the immunoprecopitated proteins mix was detected by using anti-HA antibody.
F. Schematic model of the factors involved in the maintenance of 5mC in *C. neoformans*.

Figure S3. Uhrf1 selectively binds DNA harboring hemimethylated CG dinucleotides

A. Top: Domain structure of *C. neoformans* Uhrf1 ortholog. Bottom: Coomassie Brilliant Blue staining of Uhrf1-6xHis after purification from *E. coli* and resolution by SDS-PAGE. Predicted protein domains of full-length Uhrf1 are indicated.
B. EMSA assessing competition between labeled hemimethylated DNA probe and excess concentrations unlabeled DNA (0.2, 1.0, 5.0 µM) of indicated methylation state. Where indicated, Uhrf1 protein was added at 150 nM. Graph indicates fraction of probe bound relative to condition in which no cold competitor was added; n=2, error bars represent SD.

Figure S4. Hemymethylated DNA but not unmethylated DNA is a substrate for purified Dnmt5.

A. Double stranded DNA substrates used in methyltransferase experiments. Each 20 bp substrate contains 6 CG sites that are either unmethylated, hemimethylated, or symmetrically methylated.
B. End point measurement of DNA methylation activities on methylated versus unmethylated substrates after a 6 hr reaction of 100 nM Dnmt5 and 5 μM DNA substrate. Background: no enzyme control. Error bars represent SD; n = 3.
C. DNA methylation kinetics using 30 nM Dnmt5 and 5 μM DNA substrates containing either CHG or CG motifs in an unmethylated or hemimethylated state.
D. Left: control DNA methylation reaction in which 30 nM Dnmt5 was incubated with 1 mM ATP and 5 μM of the unmethylated or hemimethylated DNA substrates. Center and right: DNA methylation reactions in which 30 nM Dnmt5 was incubated with 1 mM ATP and 5 μM unmethylated DNA in the presence or absence of 5 μM H3K9me0/3 peptide, 4 μM Uhrf1, and/or Swi6/HP1.

Figure S5. mRNA levels upon repression of *pGAL-DMT5*. The promoter region of *DMT5* was replaced with a galactose-inducible promoter (as described in Figure 3C). *pGAL-DMT5* was repressed for 10, 15, 25, 30 and 45 generations (R10-R45) and mRNA levels of *pGAL-DMT5* were compared to *pGAL-DMT5* at the starting induced condition (I_0_), wild-type cells (expressing DMT5 under its own promoter), and cell lacking the methyltransferase gene (*dmt5*Δ). RT-qPCR was normalized over *ACT1.* n*=2,* error bars represent SD.

Figure S6. WGBS of two independently-derived *RI-DMT5* strains. DNA from two independently derived strains (RI_1 and RI_2) were bisulfite treated in technical duplicate. Shown are the peaks present in both technical duplicates. Marked with asterisks are CG sites that are the symmetrically methylated and reproducible in both technical replicates.

Figure S7. Reproducibility of biological replicates for ChIP analysis.

A. Scatter plot analysis of anti-H3K9me2 and anti-FLAG ChIP. Points represent the signals for each centromere and telomere (RPKM). Each sample was compared against a biological replicate.
B. Scatter plot analysis of anti-FLAG ChIP of *RI-DMT5* compared to wild type. Pairwise comparison of the total RPKM counts for each centromere (blue) and telomere (grey). Analyses of both biological replicates are shown.

Figure S8. Stress conditions do not induce *de novo* activity in Dnmt5 To assess if specific stress conditions induce *de novo* DNA methylation activity, wild-type and RI-Dnmt5 cells were grown in the presence of stressors The genomic DNA was extracted, digested with the CG methylation sensitive endonuclease HpyCHIV and analyzed by Southern hybridization using a probe corresponding to a repetitive centromeric sequence (probe R).

Figure S9. Expression of DnmtX in *RI-DMT5*. HA-tagged version of DnmtX from *C.pinus, K. mangroviensis* and *C. bestiolae* were expressed under the control of *GAL7* promoter in an *RI-DMT5* strain. Expression of the proteins was assessed by western blotting using indicated antibodies.

Figure S10-S11. DnmtX acts as a *de novo* DNA methyltransferase in cells expressing RI-Dnmt5. BS-seq (blue) MeDIP (red) signals for centromeres and telomere (*CEN1-14, TELR/L1-14*) from cells expressing DnmtX (+DnmtX) compared to wild type and the control without DnmtX (top). Strains were analyzed after 14 days of induction of DnmtXs on YP-galactose solid media. The signals shown normalized to that of a *dmt5Δ* strain.

Figure S12. Evolutionary conservation of 5mC

A. Example of dot plot analysis of centromere alignment. Here, *CEN13* from the A1_35_8 strain was aligned to the centromeres of the other 7 species. In purple, direct alignment and in light blue, inversions.
B. Percentage of centromeric sequence covered by the pairwise alignment. Each dot in the box represents a centromere.
C. Distribution of the fragment lengths produced by pairwise alignment of centromeres. Red line: kernel density estimation (K_d_e).
D. Six families of transposable elements (Tcn1-6) were identified in the analysis. Shown is the distribution of length of the identified Tcn for each family (SE approximated by Bernoulli distribution).
E. Pearson correlations of methylation data from WGBS and Nanopore sequencing data (ONT) for the 8 strains analyzed.
F. Centromeric region of contig000014 in strain C45, as an example of 5mC calls derived from WGBS and ONT analysis. Data missing from the WGBS but present in ONT, correspond to highly repetitive regions that are not available to the WGBS pipeline.

