## Supplementary figures and images for "Evolutionary persistence of DNA methylation for millions of years after ancient loss of a *de novo* methyltransferase"

### Figure S1

*Figure S1- Catania et al.*

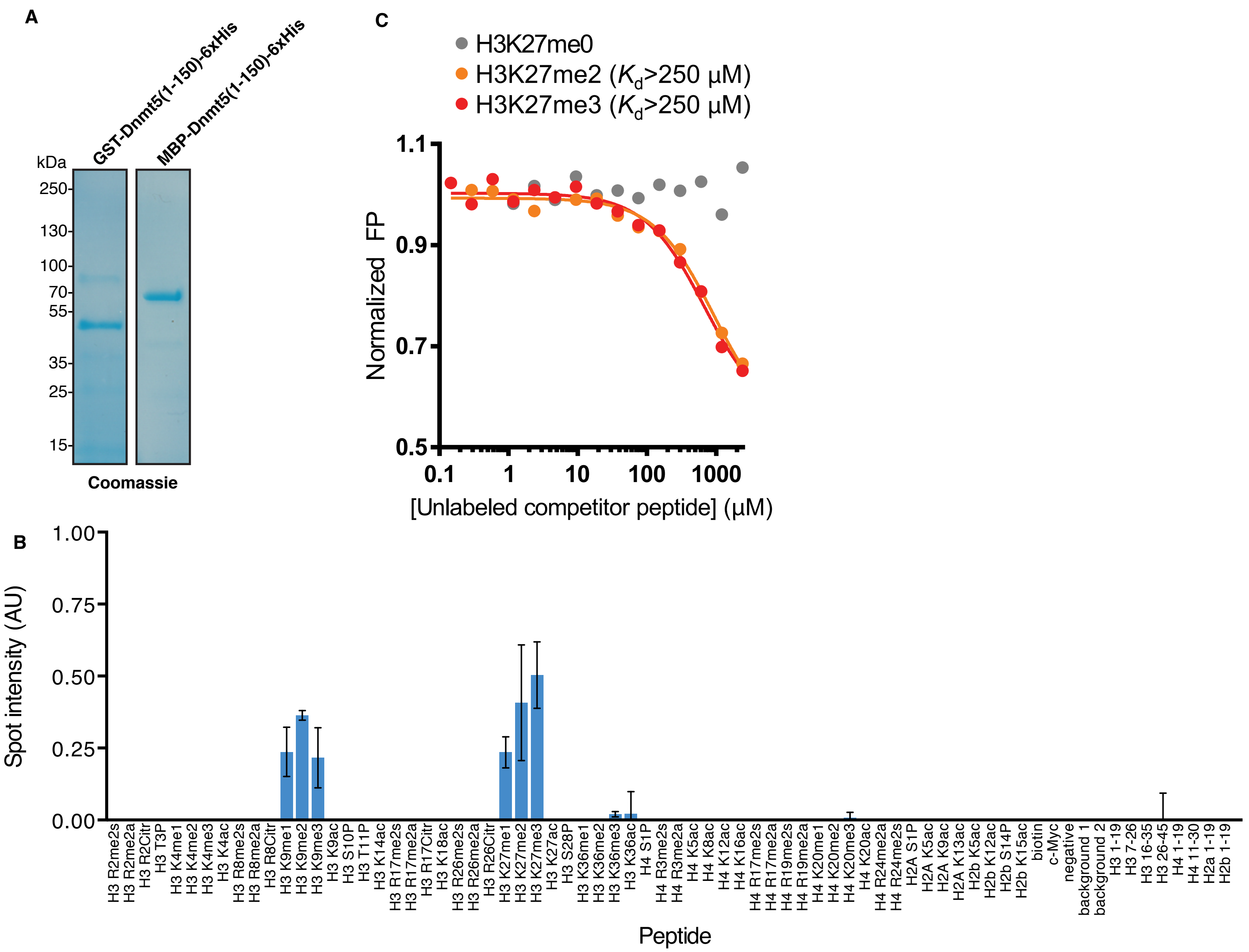

### Figure S3

*Figure S3. Catania et al.*

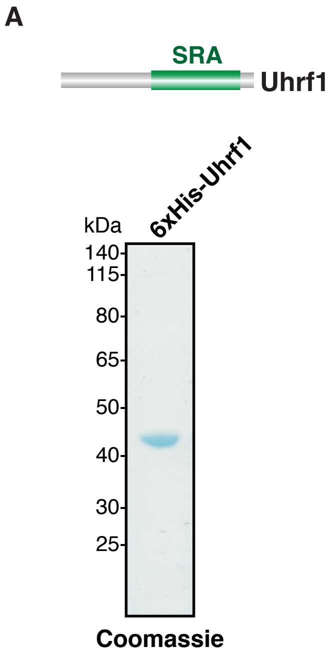


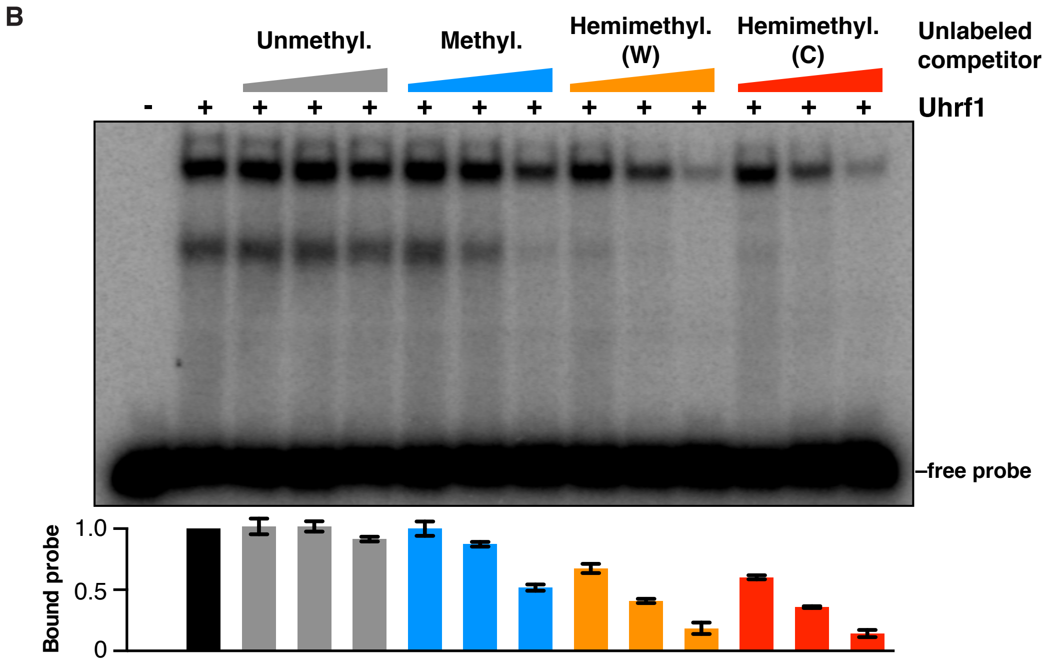

### Figure S4

*Figure S4. Catania et al.*

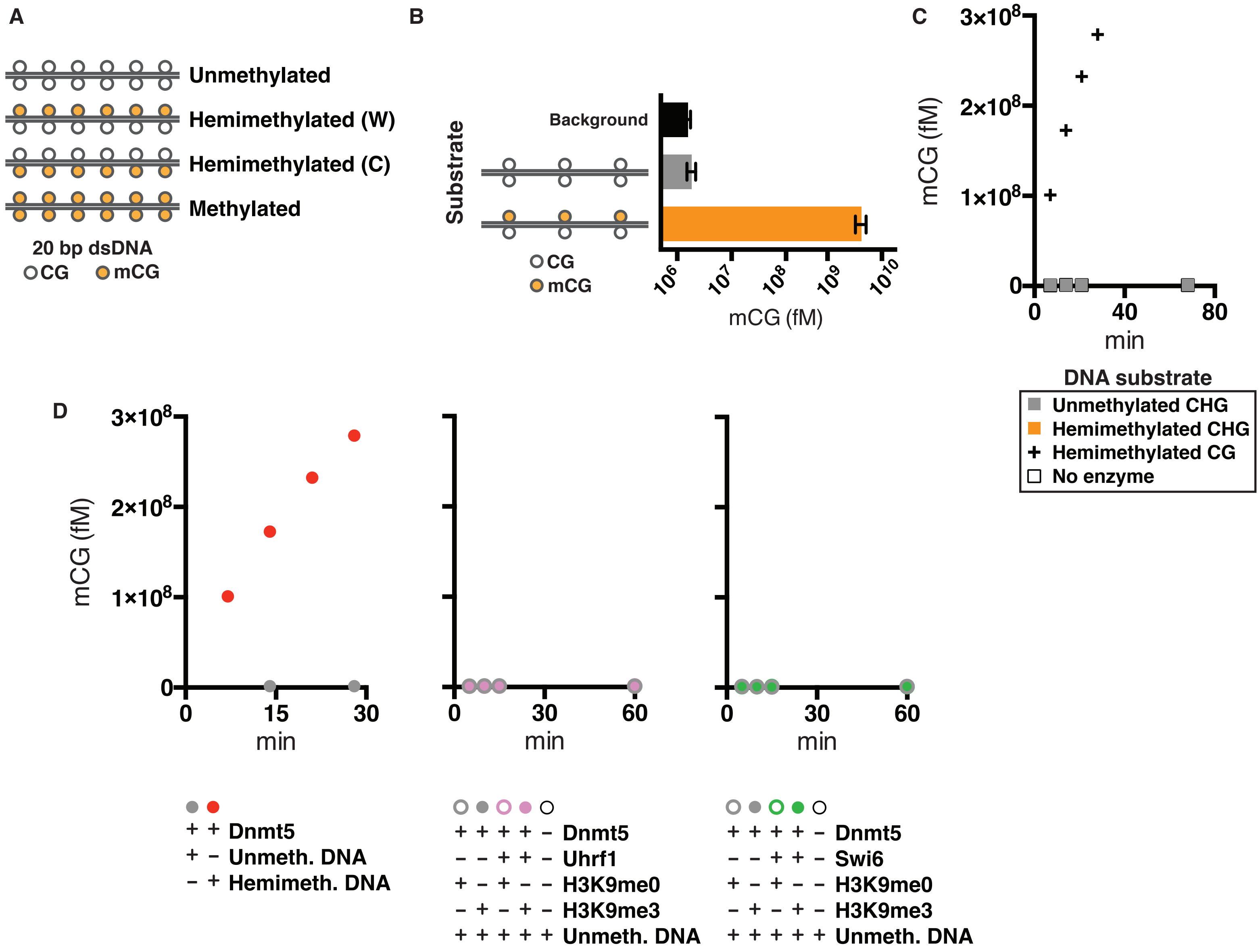

### Figure S5

***Figure S5. Catania et al.***

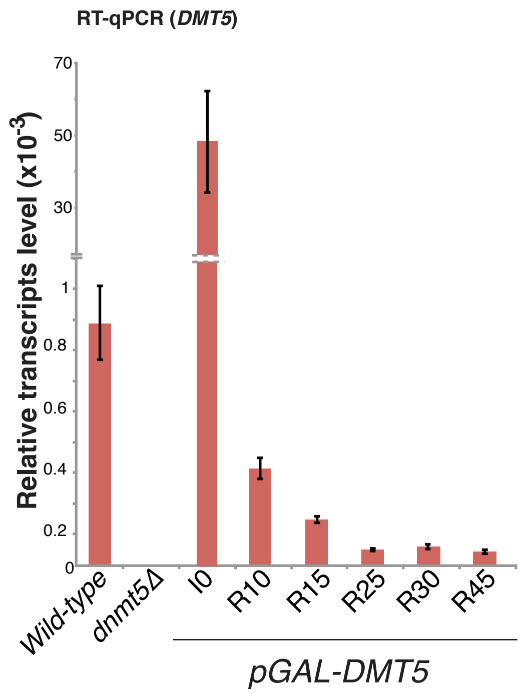

### Figure S6

*Figure S6 - Catania et al.*

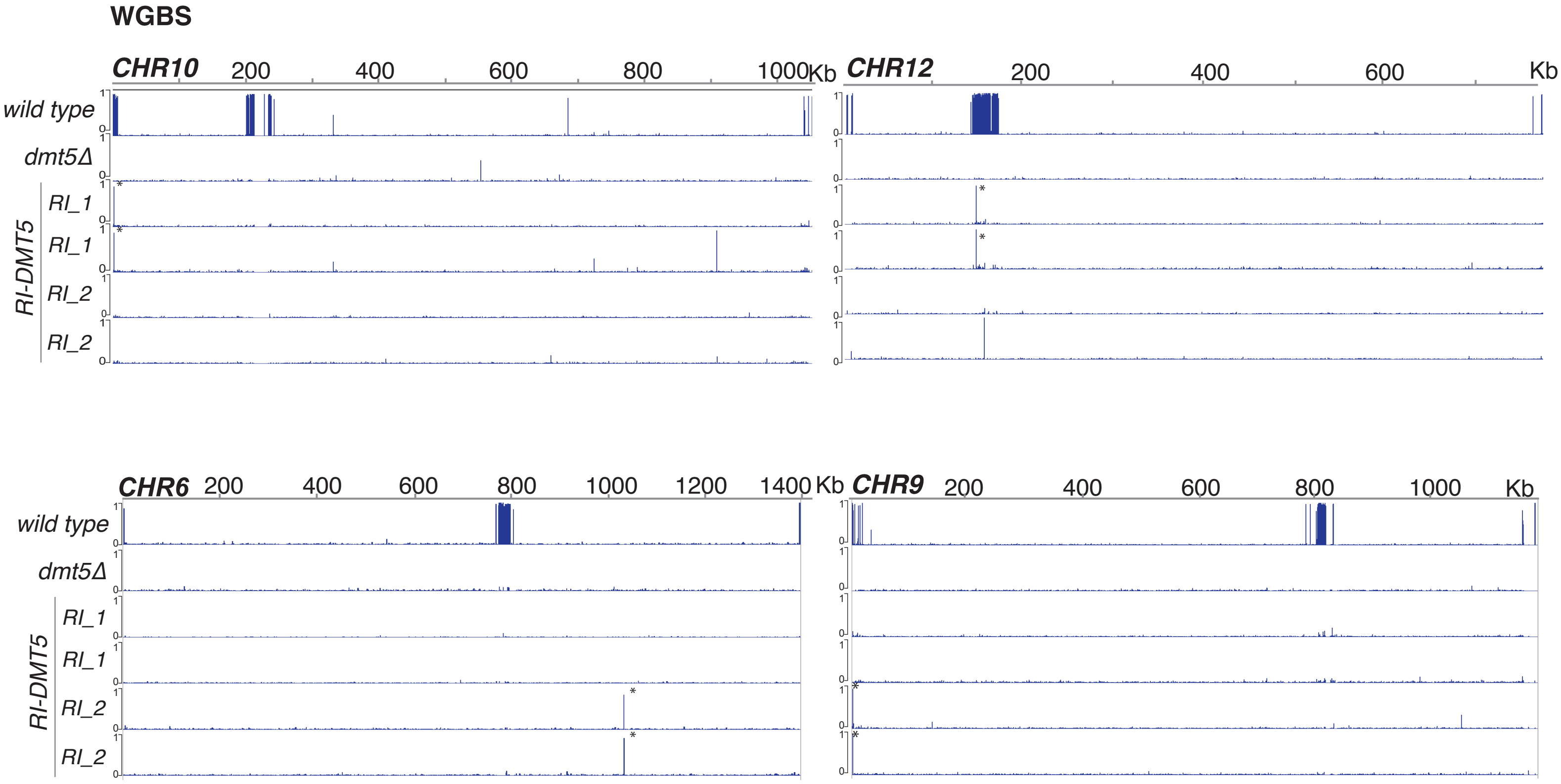

### Figure S7

*Figure S7. Catania et al.*

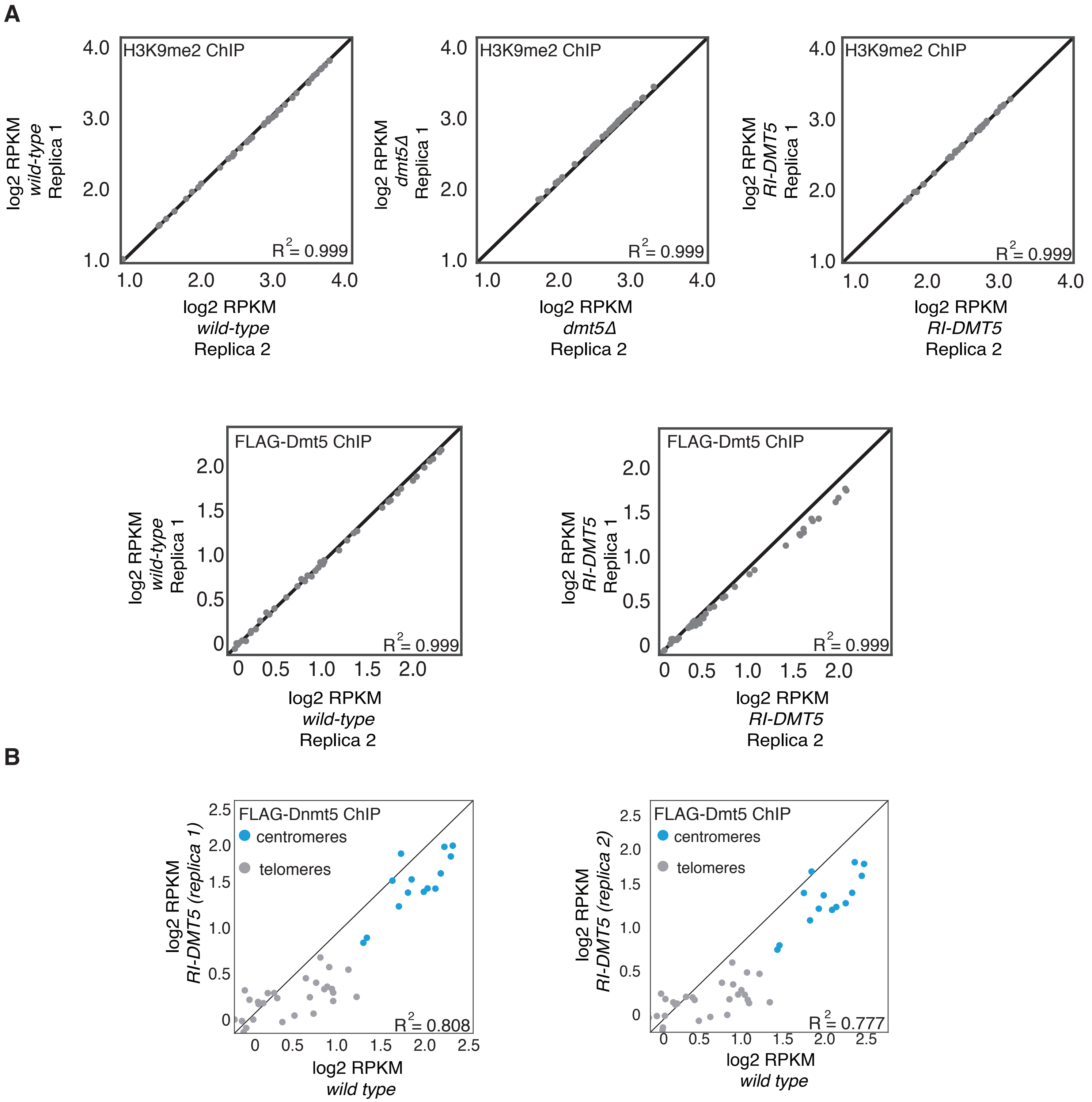

### Figure S9

***Figure S9. Catania et al.***

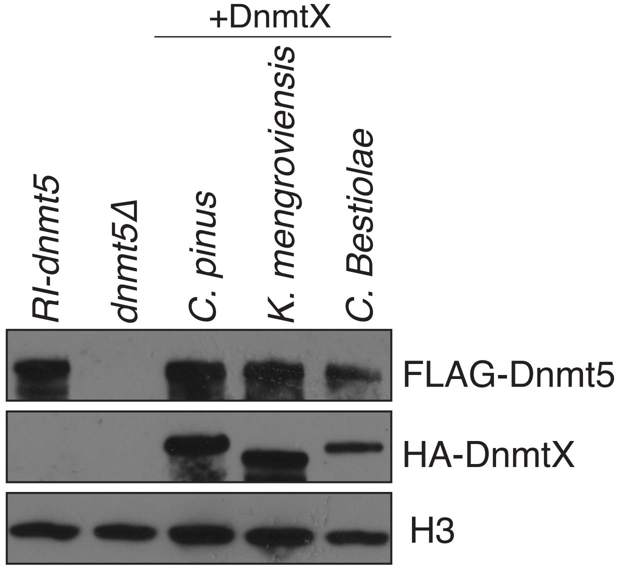

### Figure S10

##### *Figure S10. Catania et al.*

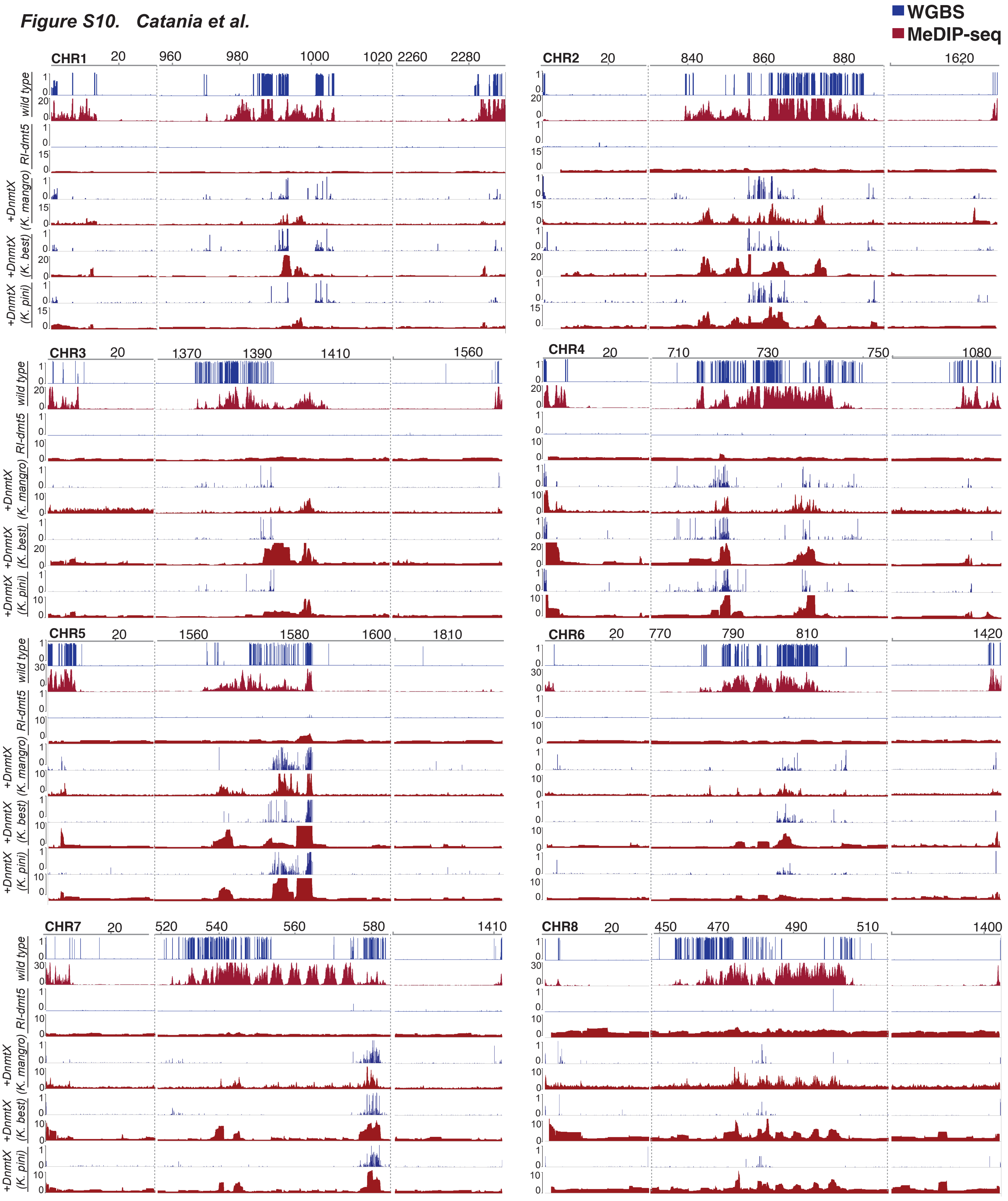

### Figure S11

#### *Figure S11. Catania et al.*

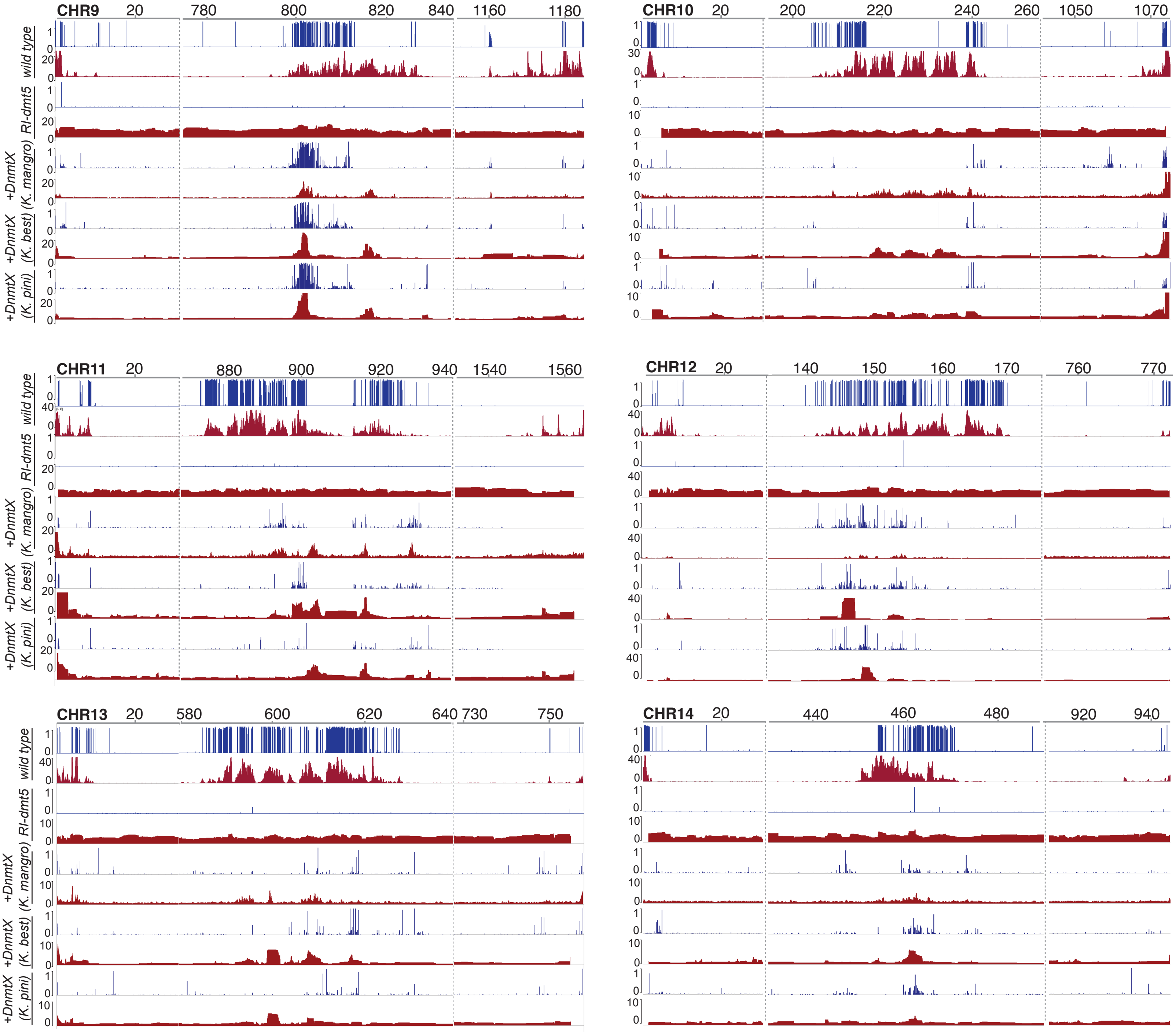

### Figure S12

*Figure S12. Catania et al.*

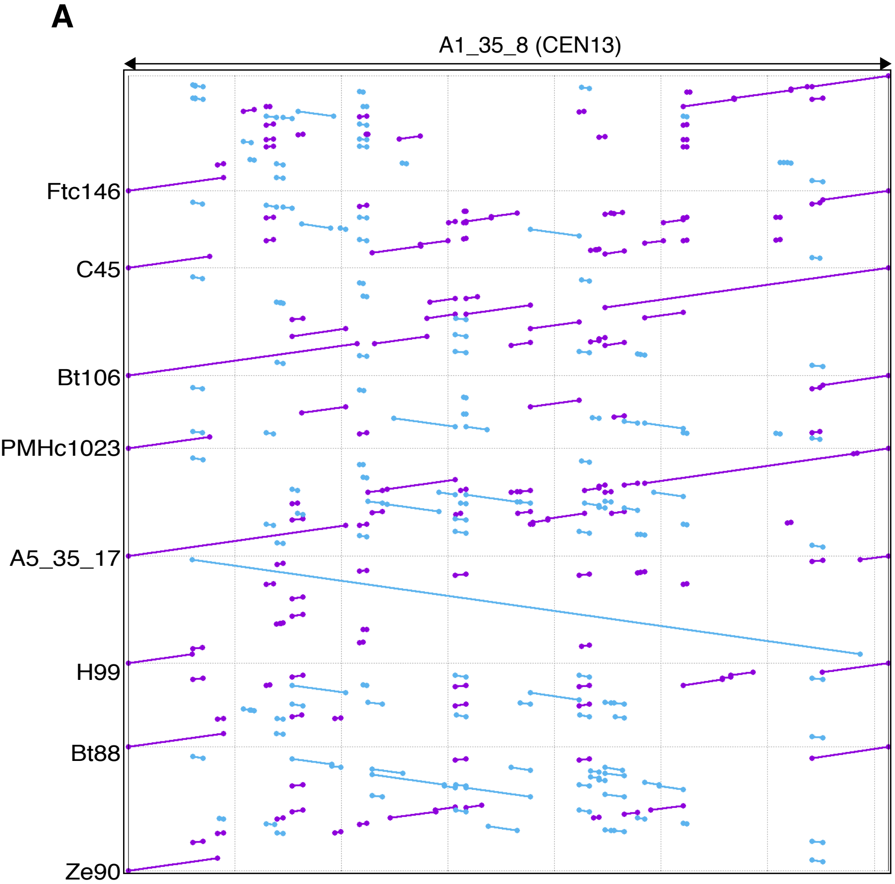


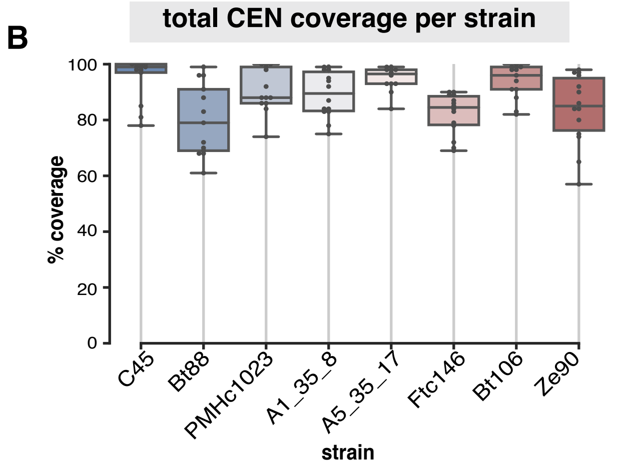


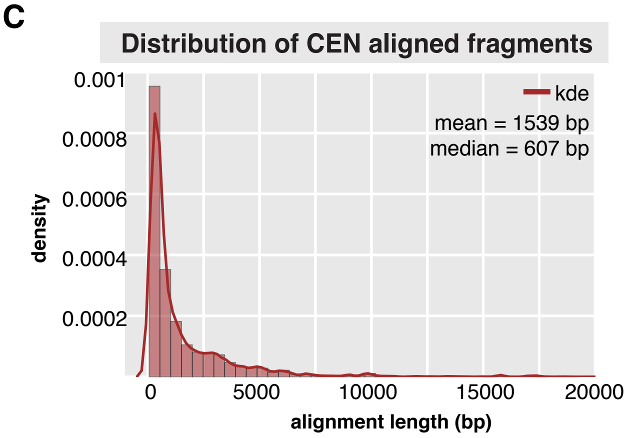


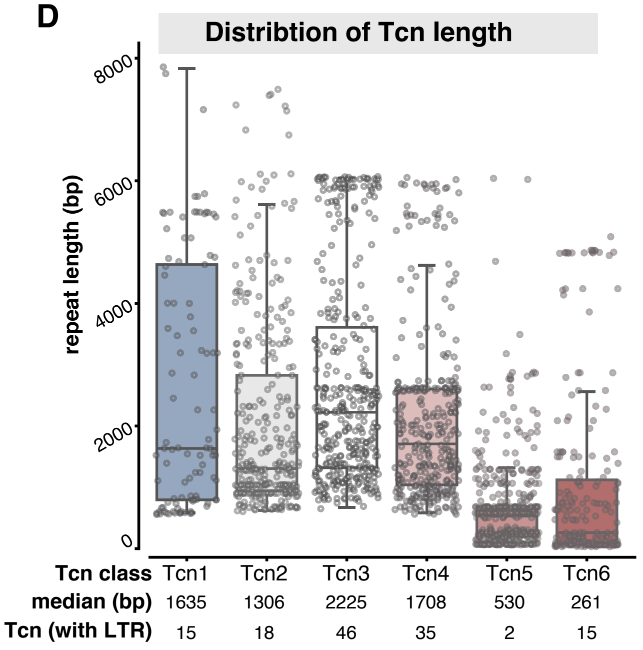


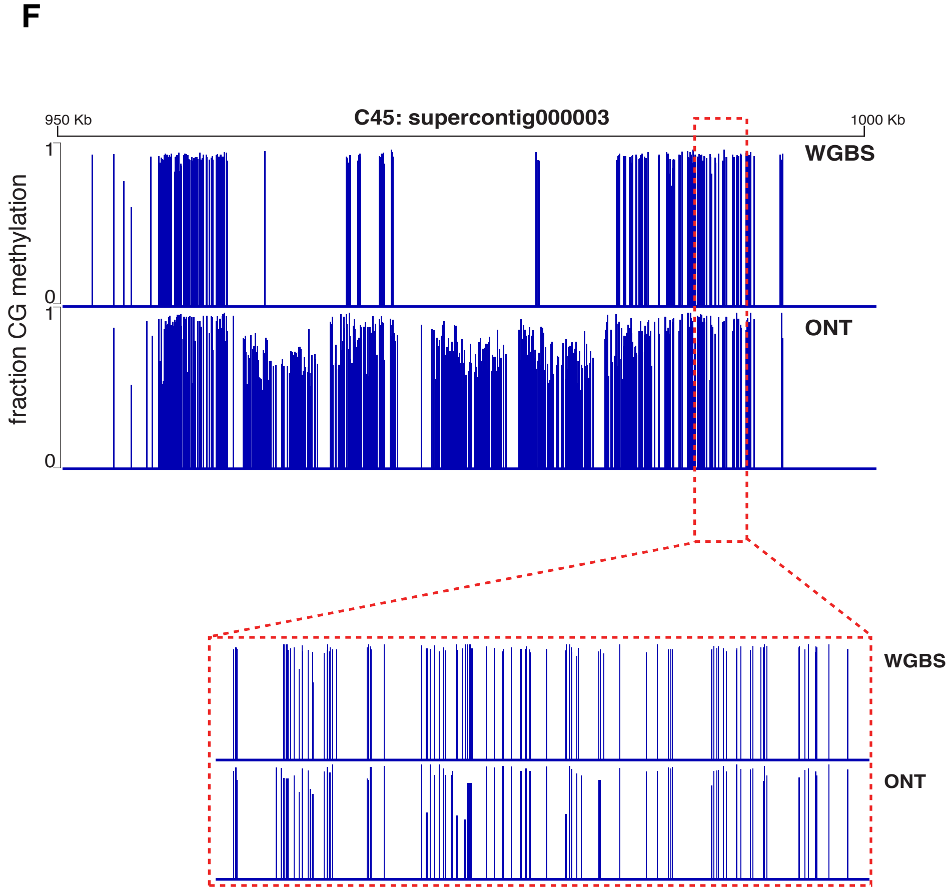


**E**

| | WGBS vs ONT |
|---|---|
| strain | Pearson coefficient |
| A1_35_8 | 0.839 |
| A5_35_17 | 0.856 |
| Bt106 | 0.894 |
| Bt88 | 0.855 |
| C45 | 0.791 |
| Ftc146 | 0.885 |
| PMHc1023 | 0.901 |
| Ze90_1 | 0.877 |

### Figures S8

***Figure S8. Catania et al.***

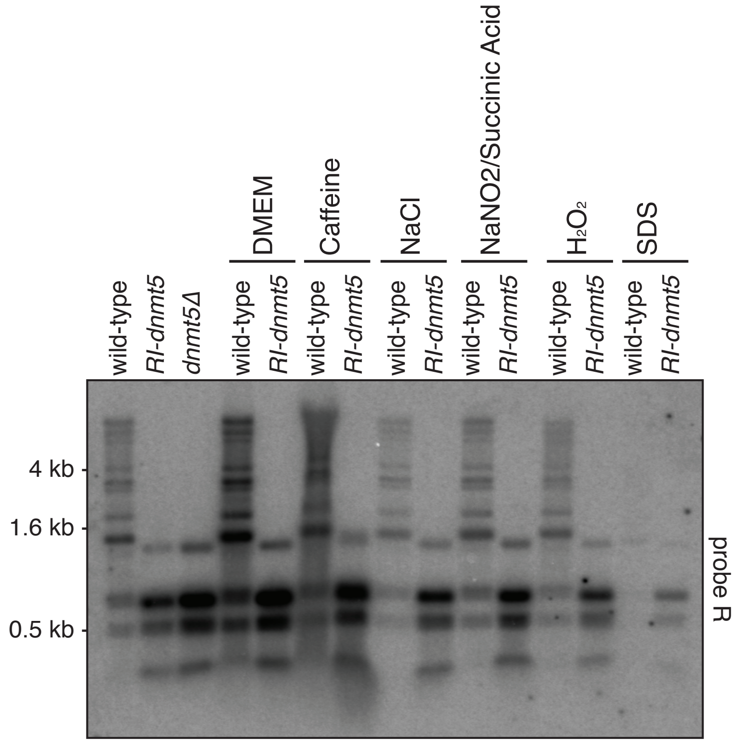
