## Supplementary material for "Evolutionary persistence of DNA methylation for millions of years after ancient loss of a *de novo* methyltransferase": Figure S2

*Figure S2.   Catania et al.*

A

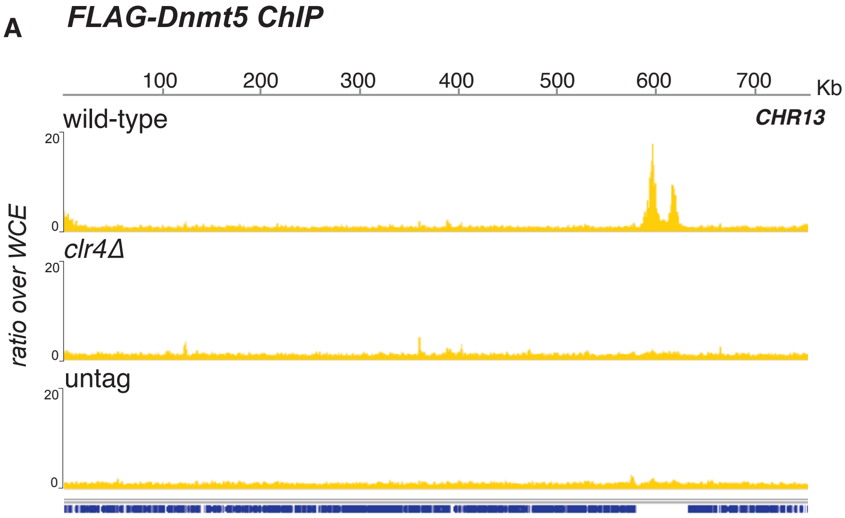


B

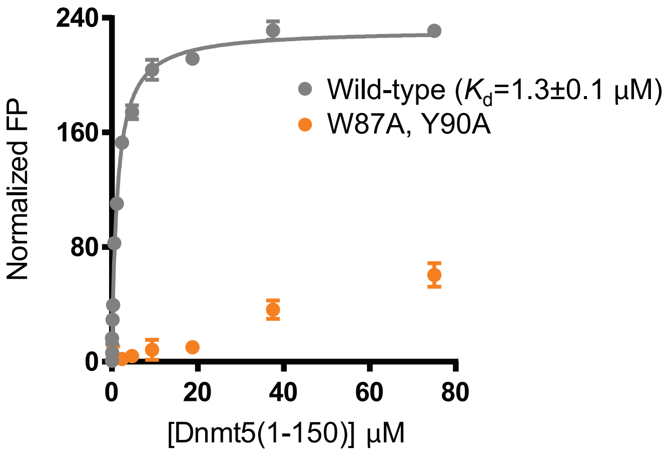


C

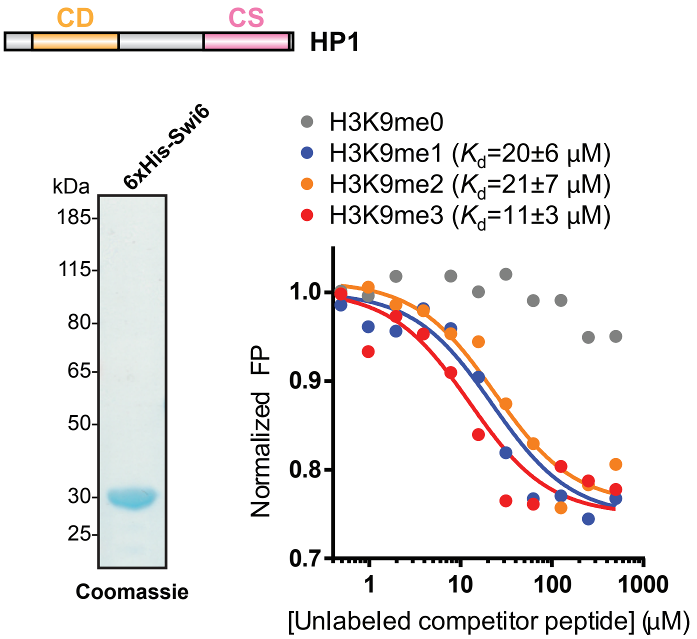


D

**Dnmt5 IP/MS**

| CNAG | peptide# | coverage | name |
|---|---|---|---|
| CNAG_07552 | 632 | 70.70% | Dmt5 |
| CNAG_03458 | 95 | 85.10% | Swi6 |
| CNAG_04828 | 36 | 84.10% | H3.3 |
| CNAG_00677 | 34 | 10.90% | Uhf1 |
| CNAG_01148 | 33 | 38.60% | Fkbp4 |
| CNAG_05391 | 28 | 26.40% | Mif2 |
| CNAG_03658 | 23 | 48.30% | |
| CNAG_08025 | 12 | 47.00% | Rnh1 |
| CNAG_00083 | 12 | 39.50% | Ccc1 |
| CNAG_07756 | 4 | 12.00% | Cdh1 |
| CNAG_01636 | 2 | 10.30% | Cpr3 |

**Swi6 IP/MS**

| CNAG | peptide# | coverage | name |
|---|---|---|---|
| CNAG_00998 | 118 | 42.10% | |
| CNAG_03458 | 56 | 56.10% | Swi6 |
| CNAG_01292 | 39 | 30.50% | Mit1 |
| CNAG_01306 | 32 | 21.70% | Mit2 |
| CNAG_01563 | 29 | 18.30% | Clr3 |
| CNAG_02922 | 18 | 18.10% | Clr2 |
| CNAG_07807 | 13 | 36.90% | H4 |
| CNAG_01648 | 13 | 36.90% | H4 |
| CNAG_07564 | 11 | 12.80% | |
| CNAG_06746 | 10 | 37.40% | H2B |
| CNAG_01406 | 9 | 40.20% | |
| CNAG_04828 | 7 | 27.50% | H3.3 |
| CNAG_06745 | 7 | 27.50% | H3 |
| CNAG_06747 | 7 | 18.30% | H2A |
| CNAG_00063 | 4 | 11.30% | Cse4 |
| CNAG_05221 | 3 | 15.90% | H2Az |

**Uhrf1 IP/MS**

| CNAG | peptide# | coverage | name |
|---|---|---|---|
| CNAG_00677 | 48 | 51% | Uhf1 |
| CNAG_04828 | 16 | 58.70% | H3.3 |

E

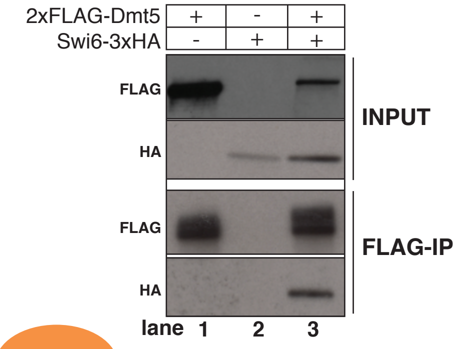


F

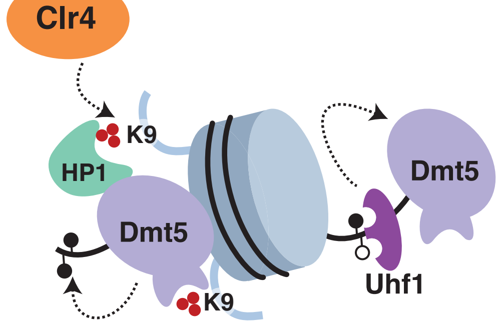
