## Supplementary material for "Evolutionary persistence of DNA methylation for millions of years after ancient loss of a *de novo* methyltransferase": Table S1

**TABLE S1: Differential methylation analysis for wild type Cryptococcus neoformans**

| **chr** | **nucleotide** | **position** | **context** | **dinucleotide context** | **methylation (g0)** | **methylation (g>120)** | **p-value** |
| --- | --- | --- | --- | --- | --- | --- | --- |
| **Colony 1** | | | | | | | |
| **Loss:** | | | | | | | |
| Chr_1 | C | 1002299 | CG | CG | 0.93 | 0 | 7.87E-09 |
| Chr_1 | G | 1002300 | CG | CG | 0.86 | 0.04 | 3.48E-07 |
| Chr_1 | C | 2288210 | CG | CG | 0.95 | 0.43 | 2.28E-02 |
| Chr_1 | G | 2288211 | CG | CG | 0.96 | 0.4 | 5.25E-03 |
| Chr_1 | C | 2288299 | CG | CG | 0.25 | 0 | 3.52E-03 |
| Chr_1 | G | 2288300 | CG | CG | 0.42 | 0 | 1.21E-06 |
| Chr_1 | C | 2288432 | CG | CG | 0.6 | 0.01 | 5.29E-11 |
| Chr_1 | G | 2288433 | CG | CG | 0.32 | 0 | 1.40E-03 |
| Chr_10 | C | 6864 | CG | CG | 0.88 | 0.01 | 3.54E-17 |
| Chr_10 | G | 6865 | CG | CG | 0.82 | 0 | 1.34E-03 |
| Chr_11 | C | 889089 | CG | CG | 0.94 | 0 | 8.92E-04 |
| Chr_11 | G | 889090 | CG | CG | 0.83 | 0 | 3.64E-09 |
| Chr_11 | C | 890047 | CG | CG | 1 | 0 | 1.91E-05 |
| Chr_11 | G | 890048 | CG | CG | 1 | 0 | 9.49E-06 |
| Chr_11 | C | 894204 | CG | CG | 1 | 0 | 1.19E-03 |
| Chr_11 | G | 894205 | CG | CG | 0.92 | 0 | 1.52E-04 |
| Chr_11 | C | 894530 | CG | CG | 1 | 0.06 | 1.26E-03 |
| Chr_11 | G | 894531 | CG | CG | 0.91 | 0 | 2.91E-07 |
| Chr_11 | G | 895890 | CG | CG | 1 | 0 | 1.20E-02 |
| Chr_11 | C | 900458 | CG | CG | 0.75 | 0.02 | 1.78E-07 |
| Chr_11 | G | 900459 | CG | CG | 0.64 | 0.05 | 4.89E-03 |
| Chr_11 | C | 922608 | CG | CG | 1 | 0.01 | 5.35E-12 |
| Chr_11 | G | 922609 | CG | CG | 0.87 | 0 | 1.64E-04 |
| Chr_11 | C | 1390056 | CG | CG | 0.48 | 0 | 9.34E-11 |
| Chr_11 | G | 1390057 | CG | CG | 0.5 | 0 | 5.99E-08 |
| Chr_12 | C | 139857 | CG | CG | 0.75 | 0 | 1.17E-07 |
| Chr_12 | G | 139858 | CG | CG | 0.91 | 0 | 2.47E-20 |
| Chr_12 | C | 153426 | CG | CG | 0.88 | 0.02 | 7.52E-09 |
| Chr_12 | G | 153427 | CG | CG | 1 | 0 | 7.13E-03 |
| Chr_12 | C | 153956 | CG | CG | 0.89 | 0.01 | 6.53E-11 |
| Chr_12 | G | 153957 | CG | CG | 0.91 | 0 | 4.66E-04 |
| Chr_12 | C | 165545 | CG | CG | 0.94 | 0 | 5.66E-09 |
| Chr_12 | G | 165546 | CG | CG | 0.94 | 0 | 4.62E-08 |
| Chr_13 | C | 589999 | CG | CG | 1 | 0 | 1.69E-02 |
| Chr_13 | G | 590000 | CG | CG | 0.93 | 0.08 | 4.18E-11 |
| Chr_13 | C | 612162 | CG | CG | 0.96 | 0 | 5.96E-03 |
| Chr_13 | G | 612163 | CG | CG | 0.97 | 0 | 2.62E-07 |
| Chr_13 | C | 612170 | CG | CG | 0.89 | 0 | 7.89E-03 |
| Chr_13 | G | 612171 | CG | CG | 0.91 | 0 | 1.60E-06 |
| Chr_13 | C | 612196 | CG | CG | 0.9 | 0 | 2.17E-03 |
| Chr_13 | G | 612197 | CG | CG | 0.88 | 0 | 2.47E-05 |
| Chr_14 | C | 567447 | CG | CG | 0.18 | 0 | 2.36E-03 |
| Chr_14 | G | 567448 | CG | CG | 0.29 | 0 | 4.04E-03 |
| Chr_2 | C | 858692 | CG | CG | 0.93 | 0.05 | 4.40E-04 |
| Chr_2 | G | 858693 | CG | CG | 0.98 | 0 | 6.23E-09 |
| Chr_2 | C | 866876 | CG | CG | 1 | 0 | 2.37E-02 |
| Chr_2 | G | 866877 | CG | CG | 0.93 | 0 | 1.12E-05 |
| Chr_3 | C | 1385539 | CG | CG | 0.96 | 0.04 | 2.13E-05 |
| Chr_3 | G | 1385540 | CG | CG | 0.97 | 0.03 | 3.76E-06 |
| Chr_3 | C | 1389958 | CG | CG | 0.94 | 0.02 | 4.01E-09 |
| Chr_3 | G | 1389959 | CG | CG | 0.93 | 0 | 2.45E-05 |
| Chr_4 | C | 720866 | CG | CG | 0.93 | 0.04 | 7.01E-05 |
| Chr_4 | G | 720867 | CG | CG | 0.93 | 0 | 3.98E-07 |
| Chr_4 | C | 721395 | CG | CG | 0.97 | 0 | 1.71E-06 |
| Chr_4 | G | 721396 | CG | CG | 0.98 | 0 | 1.94E-12 |
| Chr_4 | C | 732181 | CG | CG | 0.93 | 0 | 1.07E-06 |
| Chr_4 | G | 732182 | CG | CG | 1 | 0 | 1.12E-03 |
| Chr_4 | C | 1076440 | CG | CG | 0.89 | 0 | 8.29E-06 |
| Chr_4 | G | 1076441 | CG | CG | 0.95 | 0.01 | 3.41E-19 |
| Chr_5 | C | 1571008 | CG | CG | 0.98 | 0 | 3.81E-08 |
| Chr_5 | G | 1571009 | CG | CG | 1 | 0 | 1.03E-06 |
| Chr_5 | C | 1573858 | CG | CG | 0.9 | 0 | 2.14E-09 |
| Chr_5 | G | 1573859 | CG | CG | 0.91 | 0 | 2.13E-12 |
| Chr_5 | C | 1575127 | CG | CG | 0.93 | 0 | 4.77E-09 |
| Chr_5 | G | 1575128 | CG | CG | 0.96 | 0 | 2.01E-11 |
| Chr_6 | C | 803970 | CG | CG | 0.97 | 0 | 2.08E-08 |
| Chr_6 | G | 803971 | CG | CG | 0.93 | 0.05 | 2.00E-04 |
| Chr_6 | C | 804469 | CG | CG | 0.94 | 0 | 2.65E-12 |
| Chr_6 | G | 804470 | CG | CG | 0.94 | 0 | 1.88E-03 |
| Chr_6 | C | 805348 | CG | CG | 0.91 | 0 | 8.69E-06 |
| Chr_6 | G | 805349 | CG | CG | 0.9 | 0 | 1.15E-05 |
| Chr_6 | C | 806990 | CG | CG | 0.7 | 0 | 1.07E-09 |
| Chr_6 | G | 806991 | CG | CG | 1 | 0 | 2.85E-02 |
| Chr_6 | C | 809662 | CG | CG | 0.89 | 0 | 1.25E-05 |
| Chr_6 | G | 809663 | CG | CG | 0.98 | 0 | 1.73E-07 |
| Chr_6 | C | 811530 | CG | CG | 0.91 | 0 | 5.77E-11 |
| Chr_6 | G | 811531 | CG | CG | 0.81 | 0 | 2.32E-06 |
| Chr_7 | C | 7922 | CG | CG | 0.96 | 0 | 1.28E-05 |
| Chr_7 | G | 7923 | CG | CG | 0.89 | 0 | 2.05E-08 |
| Chr_7 | C | 539632 | CG | CG | 0.86 | 0 | 2.25E-11 |
| Chr_7 | G | 539633 | CG | CG | 0.93 | 0 | 6.15E-08 |
| Chr_7 | C | 541292 | CG | CG | 0.89 | 0 | 9.71E-03 |
| Chr_7 | G | 541293 | CG | CG | 0.92 | 0 | 1.49E-03 |
| Chr_7 | C | 578668 | CG | CG | 0.92 | 0 | 5.19E-19 |
| Chr_7 | G | 578669 | CG | CG | 0.97 | 0 | 2.19E-04 |
| Chr_8 | C | 455474 | CG | CG | 0.91 | 0.01 | 5.82E-20 |
| Chr_8 | G | 455475 | CG | CG | 1 | 0 | 1.68E-03 |
| Chr_8 | C | 1405860 | CG | CG | 0.95 | 0 | 1.49E-08 |
| Chr_8 | G | 1405861 | CG | CG | 1 | 0 | 4.38E-03 |
| Chr_9 | C | 2323 | CG | CG | 0.71 | 0 | 9.67E-08 |
| Chr_9 | G | 2324 | CG | CG | 0.51 | 0 | 2.06E-05 |
| Chr_9 | C | 808297 | CG | CG | 0.74 | 0 | 6.98E-07 |
| Chr_9 | G | 808298 | CG | CG | 0.79 | 0 | 1.24E-12 |
| Chr_9 | C | 808636 | CG | CG | 0.95 | 0 | 1.66E-07 |
| Chr_9 | G | 808637 | CG | CG | 0.96 | 0 | 8.87E-13 |
| Chr_9 | C | 812225 | CG | CG | 0.96 | 0 | 6.45E-06 |
| Chr_9 | G | 812226 | CG | CG | 0.93 | 0 | 1.97E-14 |
| Chr_9 | C | 814368 | CG | CG | 0.86 | 0.02 | 6.79E-08 |
| Chr_9 | G | 814369 | CG | CG | 0.86 | 0 | 5.44E-04 |
| Chr_9 | C | 1158332 | CG | CG | 0.88 | 0 | 5.93E-07 |
| Chr_9 | G | 1158333 | CG | CG | 0.79 | 0 | 5.34E-06 |
| **Gains:** | | | | | | | |
| Chr_2 | C | 876806 | CG | CG | 0 | 0.94 | 6.10E-08 |
| Chr_2 | G | 876807 | CG | CG | 0 | 0.91 | 3.86E-12 |
| Chr_5 | G | 1816481 | CG | CG | 0 | 0.92 | 2.32E-05 |
| Chr_5 | C | 1816480 | CG | CG | 0 | 0.98 | 3.73E-10 |
| Chr_10 | C | 5932 | CG | CG | 0 | 0.82 | 1.18E-08 |
| Chr_10 | G | 5933 | CG | CG | 0 | 0.91 | 7.21E-10 |
| Chr_9 | G | 818538 | CG | CG | 0.14 | 0.91 | 7.39E-04 |
| Chr_9 | C | 818537 | CG | CG | 0.05 | 0.96 | 7.12E-09 |
| Chr_9 | C | 810040 | CG | CG | 0.26 | 0.9 | 2.55E-03 |
| Chr_9 | G | 810041 | CG | CG | 0.25 | 0.9 | 1.10E-05 |
| **Colony 2** | | | | | | | |
| **Loss:** | | | | | | | |
| **chr** | **nucleotide** | **position** | **context** | **dinucleotide context** | **methylation (g0)** | **methylation (g>120)** | **p-value** |
| Chr_1 | C | 992240 | CG | CG | 0.83 | 0 | 3.58E-02 |
| Chr_1 | G | 992241 | CG | CG | 0.96 | 0.24 | 2.73E-04 |
| Chr_1 | C | 1002101 | CG | CG | 0.95 | 0.45 | 2.64E-02 |
| Chr_1 | G | 1002102 | CG | CG | 1 | 0.3 | 5.84E-03 |
| Chr_1 | C | 1003458 | CG | CG | 0.97 | 0.26 | 1.98E-05 |
| Chr_1 | G | 1003459 | CG | CG | 0.93 | 0.29 | 1.57E-03 |
| Chr_10 | C | 2930 | CG | CG | 0.92 | 0.27 | 1.54E-03 |
| Chr_10 | G | 2931 | CG | CG | 0.98 | 0.33 | 1.40E-03 |
| Chr_10 | C | 216082 | CG | CG | 0.98 | 0.36 | 3.33E-03 |
| Chr_10 | G | 216083 | CG | CG | 0.92 | 0.38 | 5.32E-02 |
| Chr_11 | C | 897688 | CG | CG | 0.94 | 0.22 | 6.24E-03 |
| Chr_11 | G | 897689 | CG | CG | 0.92 | 0.3 | 7.81E-04 |
| Chr_11 | C | 898323 | CG | CG | 0.98 | 0.31 | 5.97E-04 |
| Chr_11 | G | 898324 | CG | CG | 0.98 | 0.26 | 9.51E-04 |
| Chr_11 | C | 918662 | CG | CG | 0.95 | 0.2 | 6.59E-05 |
| Chr_11 | G | 918663 | CG | CG | 0.95 | 0.37 | 2.34E-02 |
| Chr_11 | C | 921554 | CG | CG | 0.93 | 0.24 | 3.51E-04 |
| Chr_11 | G | 921555 | CG | CG | 0.96 | 0.37 | 4.99E-03 |
| Chr_12 | C | 139857 | CG | CG | 0.75 | 0 | 8.45E-09 |
| Chr_12 | G | 139858 | CG | CG | 0.91 | 0 | 2.93E-12 |
| Chr_12 | C | 153282 | CG | CG | 0.82 | 0.29 | 1.07E-03 |
| Chr_12 | G | 153283 | CG | CG | 0.78 | 0.36 | 3.41E-02 |
| Chr_13 | C | 598201 | CG | CG | 1 | 0.24 | 1.33E-02 |
| Chr_13 | G | 598202 | CG | CG | 1 | 0.21 | 2.56E-03 |
| Chr_13 | C | 611396 | CG | CG | 0.93 | 0.47 | 6.34E-02 |
| Chr_13 | G | 611397 | CG | CG | 0.98 | 0.24 | 2.22E-03 |
| Chr_13 | C | 627160 | CG | CG | 0.93 | 0.32 | 1.00E-02 |
| Chr_13 | G | 627161 | CG | CG | 0.98 | 0.27 | 2.22E-03 |
| Chr_14 | C | 2714 | CG | CG | 0.96 | 0.25 | 2.19E-04 |
| Chr_14 | G | 2715 | CG | CG | 0.91 | 0.36 | 1.59E-02 |
| Chr_14 | C | 462659 | CG | CG | 0.91 | 0.27 | 3.32E-02 |
| Chr_14 | G | 462660 | CG | CG | 0.96 | 0.22 | 5.71E-03 |
| Chr_14 | C | 467569 | CG | CG | 1 | 0.32 | 4.54E-02 |
| Chr_14 | G | 467570 | CG | CG | 0.91 | 0.31 | 1.58E-02 |
| Chr_4 | C | 710494 | CG | CG | 0.98 | 0.33 | 9.30E-04 |
| Chr_4 | G | 710495 | CG | CG | 0.94 | 0.25 | 3.65E-04 |
| Chr_5 | C | 1802702 | CG | CG | 0.94 | 0.31 | 1.05E-03 |
| Chr_5 | G | 1802703 | CG | CG | 0.92 | 0.28 | 8.53E-04 |
| Chr_6 | C | 804817 | CG | CG | 0.89 | 0 | 1.68E-05 |
| Chr_6 | G | 804818 | CG | CG | 0.96 | 0 | 8.85E-07 |
| Chr_6 | C | 805554 | CG | CG | 0.95 | 0.28 | 4.83E-03 |
| Chr_6 | G | 805555 | CG | CG | 0.95 | 0.29 | 1.49E-03 |
| Chr_6 | C | 810462 | CG | CG | 1 | 0.24 | 4.67E-04 |
| Chr_6 | G | 810463 | CG | CG | 0.96 | 0.19 | 2.70E-04 |
| Chr_6 | C | 811705 | CG | CG | 1 | 0.21 | 1.90E-02 |
| Chr_6 | G | 811706 | CG | CG | 0.9 | 0.38 | 5.97E-03 |
| Chr_6 | C | 819783 | CG | CG | 0.92 | 0.29 | 1.79E-03 |
| Chr_6 | G | 819784 | CG | CG | 0.87 | 0.25 | 1.14E-05 |
| Chr_7 | C | 537058 | CG | CG | 0.95 | 0.28 | 4.74E-03 |
| Chr_7 | G | 537059 | CG | CG | 0.93 | 0.17 | 3.12E-05 |
| Chr_7 | C | 537071 | CG | CG | 0.95 | 0.21 | 1.40E-03 |
| Chr_7 | G | 537072 | CG | CG | 1 | 0.17 | 4.37E-06 |
| Chr_8 | C | 463076 | CG | CG | 0.93 | 0 | 1.54E-05 |
| Chr_8 | G | 463077 | CG | CG | 0.95 | 0 | 4.79E-11 |
| Chr_8 | C | 467094 | CG | CG | 1 | 0.21 | 2.36E-02 |
| Chr_8 | G | 467095 | CG | CG | 0.94 | 0.27 | 2.54E-04 |
| Chr_8 | C | 471300 | CG | CG | 0.95 | 0.23 | 1.26E-04 |
| Chr_8 | G | 471301 | CG | CG | 0.95 | 0.35 | 2.39E-02 |
| Chr_8 | C | 478918 | CG | CG | 0.97 | 0.31 | 1.20E-02 |
| Chr_8 | G | 478919 | CG | CG | 0.93 | 0.19 | 2.33E-02 |
| Chr_9 | C | 807958 | CG | CG | 0.91 | 0.38 | 5.60E-02 |
| Chr_9 | G | 807959 | CG | CG | 0.92 | 0.29 | 3.90E-04 |
| Chr_9 | C | 808775 | CG | CG | 0.95 | 0.25 | 8.35E-04 |
| Chr_9 | G | 808776 | CG | CG | 0.96 | 0.35 | 7.48E-04 |
| Chr_9 | C | 811455 | CG | CG | 0.92 | 0.28 | 2.77E-03 |
| Chr_9 | G | 811456 | CG | CG | 0.97 | 0.25 | 3.82E-05 |
| Chr_9 | C | 814774 | CG | CG | 0.81 | 0.19 | 3.90E-04 |
| Chr_9 | G | 814775 | CG | CG | 0.85 | 0.33 | 6.91E-03 |
| Chr_9 | C | 814968 | CG | CG | 0.85 | 0.28 | 3.15E-03 |
| Chr_9 | G | 814969 | CG | CG | 0.74 | 0.12 | 8.65E-05 |
| Chr_9 | C | 815499 | CG | CG | 0.83 | 0.34 | 3.22E-04 |
| Chr_9 | G | 815500 | CG | CG | 0.76 | 0.3 | 5.47E-03 |
| Chr_9 | C | 818527 | CG | CG | 0.93 | 0.25 | 6.87E-05 |
| Chr_9 | G | 818528 | CG | CG | 0.97 | 0.24 | 9.47E-03 |
| Chr_9 | C | 1158332 | CG | CG | 0.88 | 0.25 | 2.69E-04 |
| Chr_9 | G | 1158333 | CG | CG | 0.79 | 0.36 | 3.53E-02 |
| **Gains:** | | | | | | | |
| Chr_13 | G | 18604 | CG | CG | 0 | 0.67 | 5.92E-08 |
| Chr_13 | C | 18603 | CG | CG | 0 | 0.67 | 2.10E-06 |
| Chr_13 | G | 753513 | CG | CG | 0.06 | 0.59 | 5.46E-06 |
| Chr_13 | C | 753512 | CG | CG | 0.05 | 0.54 | 3.60E-04 |
| Chr_2 | C | 878899 | CG | CG | 0 | 0.77 | 9.78E-08 |
| Chr_2 | G | 878900 | CG | CG | 0 | 0.74 | 4.98E-06 |
| Chr_5 | C | 7078 | CG | CG | 0 | 0.75 | 1.84E-07 |
| Chr_5 | G | 7079 | CG | CG | 0 | 0.54 | 3.85E-07 |
| Chr_7 | C | 551115 | CG | CG | 0 | 0.64 | 6.54E-04 |
| Chr_7 | G | 551116 | CG | CG | 0 | 0.62 | 5.29E-03 |
