## Supplementary material for "Evolutionary persistence of DNA methylation for millions of years after ancient loss of a *de novo* methyltransferase": Table S2

| TABLE S2. List of strains used in this study | | | | |
| --- | --- | --- | --- | --- |
| Strain number | **Genotype** | **Parent** | ***Notes*** | **Fig.** |
| CM229 | Wild type | CM018 | Wild type | 1 |
| CM1197 | *CNAG_05404∆::natR* | CM229 | *clr4∆* | 1 |
| CM1837 | *CNAG_07552∆::kanR* | CM229 | *dmt5∆-*1.5 kb from ATG | 1 |
| CM1838 | *CNAG_00677∆::natR* | CM229 | *uhf1∆* | 1 |
| CM1839 | *CNAG_05404∆::natR CNAG_00677∆::kanR* | CM1197 | *clr4∆ uhf1∆* | 1 |
| CM1840 | *hygR-2xFLAG-CNAG_07552-W87A Y90A* | CM229 | *CDmut-dmt5* | 1 |
| CM1841 | *CNAG_03458∆::kanR* | CM229 | *swi6∆* | 1 |
| CM1842 | *hygR::2xFLAG-CNAG_07552-W87A Y90A CNAG_03458∆::kanR* | CM1840 | *CDmut-dmt5 swi6∆* | 1 |
| CM1843 | *hygR::2xFLAG-CNAG_07552* | CM229 | *FLAG-DMT5* | 1,3,4 |
| CM1844 | *hygR::pGAL7-2xFLAG-CNAG_07552* | CM229 | *pGAL-DMT5-OFF* | 3A |
| CM1845 | *hygR::pGAL7-2xFLAG-CNAG_07552* | CM229 | *pGAL-DMT5-ON* | 3B-C |
| CM1846 | *hygR::2xFLAG-CNAG_07552-RI* | CM1837 | *RI-DMT5* | 4 |
| CM1847 | *hygR::2xFLAG-CNAG_07552-RI* | KN99 | *RI-DMT5* | 5 |
| CM1848 | *MATa CEN13:natR* | KN99a |  | 4E |
| CM1849 | *MATα CEN2:kanR hygR-2xFLAG-CNAG_07552-RI* | KN99α |  | 4E |
| CM1850 | *CEN13::natR CEN2::kanR hygR-2xFLAG- CNAG_07552-RI* | KN99 | CM1848x1849 | 4E |
| CM1851 | *CEN13::natR CEN2::kanR hygR-2xFLAG- CNAG_07552-RI* | KN99 | CM1848x1849 | 4E |
| CM1852 | *CEN13::natR CEN2::kanR hygR-2xFLAG- CNAG_07552-RI* | KN99 | CM1848x1849 | 4E |
| CM1853 | *CEN13::natR CEN2::kanR hygR-2xFLAG-CNAG_07552-RI* | KN99 | CM1848x1849 | 4E |
| CM1854 | *CEN13::natR CEN2::kanR hygR-2xFLAG- CNAG_07552-RI* | KN99 | CM1848x1849 | 4E |
| CM1855 | *CEN13::natR CEN2::kanR hygR-2xFLAG- CNAG_07552-RI* | KN99 | CM1848x1849 | 4E |
| CM1856 | *CEN13::natR CEN2::kanR hygR-2xFLAG- CNAG_07552-RI* | KN99 | CM1848x1849 | 4E |
| CM1857 | *CEN13::natR CEN2::kanR hygR-2xFLAG- CNAG_07552-RI* | KN99 | CM1848x1849 | 4E |
| CM1858 | *CEN13:natR CEN2::kanR hygR-2xFLAG- CNAG_07552-RI* | KN99 | CM1848x1849 | 4E |
| CM1859 | *CEN13:natR CEN2::kanR hygR-2xFLAG- CNAG_07552-RI* | KN99 | CM1848x1849 | 4E |
| CM1860 | *MATα CEN13::natR* | KN99α |  | 4F |
| CM1861 | *MATa CEN2::hygR CNAG_07552∆::kanR* | KN99a |  | 4F |
| CM1862 | *CEN13:natR CEN2:hygR* | KN99 | CM1861x CM1860 | 4F |
| CM1863 | *CEN13::natR CEN2:hygR* | KN99 | CM1861x CM1860 | 4F |
| CM1864 | *CEN13::natR CEN2::hygR* | KN99 | CM1861x CM1860 | 4F |
| CM1865 | *CEN13::natR CEN2:hygR CNAG_07552∆::kanR* | KN99 | CM1861x CM1860 | 4F |
| CM1866 | *MATα CEN2::hygR* | KN99α |  | 4G |
| CM1867 | *MATa CEN13::natR CNAG_07552∆::kanR* | KN99a |  | 4G |
| CM1868 | *CEN13::natR CEN2::hygR* | KN99 | CM1866 x CM1867 | 4G |
| CM1869 | *CEN13::natR CEN2::hygR* | KN99 | CM1866 x CM1867 | 4G |
| CM1870 | *CEN13::natR CEN2::hygR* | KN99 | CM1866 x CM1867 | 4G |
| CM1871 | *CEN13::natR CEN2::hygR CNAG_07552∆::* kanR | KN99 | CM1866 x CM1867 | 4G |
| CM1872 | *ura5∆::pGAL7-3xHA-I203_05465-natR hygR-2xFLAG- CNAG_07552-RI* | KN99 | DnmtX from *K. mengroviensis* | 5A-B |
| CM1873 | *ura5∆::pGAL7-3xHA-I206_02856-natR hygR-2xFLAG- CNAG_07552-RI* | KN99 | DnmtX from *C. pinus* | 6A-B |
| CM1874 | *ura5∆::pGAL7-3xHA-I302_00877-natR hygR-2xFLAG- CNAG_07552-RI* | KN99 | DnmtX from *C. bestiolae* | 6A-B |
| CM1910 | *CNAG_03458-3xHA:kanR* | CM229 | Swi6-HA | S2 |
| CM1911 | *CNAG_3458-3xHA:: kanR 2xFlag -CNAG_07552::hygR* | CM1843 | Swi6-HA Flag-Dnmt5 | S2 |
| CM1912 | *CNAG_07552-CBP-2xFlag::natR* | CM229 | Dnmt5-CBP-FLAG | S2 |
| CM1913 | *CNAG_03458-2xFLAG-CBP::natR* | CM229 | Swi6-CBP-FLAG | S2 |
| CM1914 | *CNAG_00677-2xFLAG-CBP::natR* | CM229 | Uhf1-CBP-FLAG | S2 |
| CM1915 | *CEN4:Fragment-kanR* | CM229 | control | 5 |
| CM1916 | CEN4*:*:Fragment-::kanR | CM229 | control | 5 |
| CM1917 | *CEN4::Fragment+::kanR-Hpa*II*-METHYLATED* | CM229 | methylated | 5 |
| CM1918 | *CEN4::Fragment::kanR-Hpa*II*-METHYLATED* | CM229 | methylated | 5 |
| CM1919 | *CEN4::Fragment:: kanR -Hpa*II*-METHYLATED* | CM229 | methylated | 5 |
| CM1920 | *CEN4::Fragment:: kanR -Hpa*II*-METHYLATED* | CM229 | methylated | 5 |
| CM1921 | Wild type | CM229 | g0 | 6C-E |
| CM1922 | propagated wild-type | CM229 | colony 1 | 6C-E |
| CM1923 | propagated wild-type | CM229 | colony 2 | 6C-E |
| CM1875 | C45 | JPC123 | gift from Perfect Lab | 7 |
| CM1876 | A1_35_8 | JPC160 | gift from Perfect Lab | 7 |
| CM1877 | A5_35_17 | JPC161 | gift from Perfect Lab | 7 |
| CM1878 | Bt106 | JPC123 | gift from Perfect Lab | 7 |
| CM1879 | Bt88 | JPC54 | gift from Perfect Lab | 7 |
| CM1880 | Ze90_1 | JPC20 | gift from Perfect Lab | 7 |
| CM1882 | FTC146-1 | JPC260 | gift from Perfect Lab | 7 |
| CM1883 | PMHc1023.ENR | JPC327 | gift from Perfect Lab | 7 |
