## Supplementary material for "Evolutionary persistence of DNA methylation for millions of years after ancient loss of a *de novo* methyltransferase": Table S3

**TABLE S3. Oligonucletides used in this study**

| C# | Sequence (M=5mC) | Notes | Strand | Assay |
| --- | --- | --- | --- | --- |
| 6153 | CCCACTGGATGAAACCTCGT | Probe U/cen13 | 5’ primer | Southern |
| 6154 | TCGGTGGGATGAGCAGAAAAAC | ProbeU/cen13 | 3’ primer |  |
| 6155 | TGGACCGGAACACCGTAGA | Probe R | 5’ primer | Southern |
| 6156 | ATACCGGTACGGGTGCATG | Probe R | 3’ primer |  |
| 6665 | CACGCGACGCACGACGCGAA | Unmethylated | Watson | EMSA, DMT |
| 6664 | TTCGCGTCGTGCGTCGCGTG |  | Crick |  |
| 6661 | CAMGMGAMGCAMGAMGMGAA | Hemimethylated (W) | Watson | EMSA, DMT |
| 6664 | TTCGCGTCGTGCGTCGCGTG |  | Crick |  |
| 6665 | CACGCGACGCACGACGCGAA | Hemimethylated (C) | Watson | EMSA, DMT |
| 6662 | TTMGMGTMGTGMGTMGMGTG |  | Crick |  |
| 6661 | CAMGMGAMGCAMGAMGMGAA | Methylated | Watson | EMSA, DMT |
| 6662 | TTMGMGTMGTGMGTMGMGTG |  | Crick |  |
| 7289 | CATGGCCTAAGCCGGACTGAATGAGCAAGCTTCAGGAGAATTCTGCCGGACTGCAGATGC | A | Watson | DMT |
| 7290 | GCATCTGCAGTCCGGCAGAATTCTCCTGAAGCTTGCTCATTCAGTCCGGCTTAGGCCATG |  | Crick |  |
| 7287 | CATGGCCTAAGCCGGACTGAATGAGCAAGCTTCMGGAGAATTCTGCCGGACTGCAGATGC | B | Watson | DMT |
| 7288 | GCATCTGCAGTCCGGCAGAATTCTCMGGAAGCTTGCTCATTCAGTCCGGCTTAGGCCATG |  | Crick |  |
| 7291 | CATGGCCTAAGCAGGACTGAATGAGCAAGCTTCAGGAGAATTCTGCAGGACTGCAGATGC | C | Watson | DMT |
| 7292 | GCATCTGCAGTCCTGCAGAATTCTCCTGAAGCTTGCTCATTCAGTCCTGCTTAGGCCATG |  | Crick |  |
| 7777 | CATGGCCTAAGCAGGACTGAATGAGCAAGCTTCMAGAGAATTCTGCAGGACTGCAGATGC | Hemimethylated CHG | Watson | DMT |
| 7896 | GCATCTGCAGTCCTGCAGAATTCTCTGGAAGCTTGCTCATTCAGTCCTGCTTAGGCCATG |  | Crick |  |
| 7817 | CATGGCCTAAGCAGGACTGAATGAGCAAGCTTCCAGAGAATTCTGCAGGACTGCAGATGC | Unmethylated CHG | Watson | DMT |
| 7896 | GCATCTGCAGTCCTGCAGAATTCTCTGGAAGCTTGCTCATTCAGTCCTGCTTAGGCCATG |  | Crick |  |
| 8235 | CAGCATCGTTTGAGTAAGGC | qDMT5_F | 5’ primer | qPCR |
| 8236 | AGCTTTGGAAATGCAAGCACGC | qDMT5_F | 3’ primer |  |
| 8359 | TATCCTCCGTATCGATCTTGC | qactF1 | 5’ primer | qPCR |
| 8360 | AAGCTTCTCCTTAATGTCTC | qactR1 | 3’ primer |  |
| 8346 | GGAGAATGGAGATGAGGTGAAAAGGG |  | 5’ primer | qPCR (1) |
| 8347 | GTACGAGACGACCACGAAGC |  | 3’ primer |  |
| 8348 | TTCGTCGCGTACGGGGACG |  | 5’ primer | qPCR(2) |
| 8349 | GGCGACCTCGATGTCCTC |  | 3’ primer |  |
| 8350 | GACCGTCGAGGACATCGAG |  | 5’ primer | qPCR (3) |
| 8351 | CCCCGCTCGCGGGCGAACT |  | 3’ primer |  |
| 8352 | GTCGGGCGCGCGTTGATG |  | 5’ primer | qPCR (4) |
| 8353 | TGCGTTAACGTTGGTGACC |  | 3’ primer |  |
| 8354 | ACCTCTGGCTGGAGGTCAC |  | 5’ primer | qPCR (5) |
|  | CCTTCTCCTACCCATTTCCCTTTCCTTACC |  | 3’ primer |  |
| 8355 | AACAGACAATCGGCTGCTCTG |  | 5’ primer | qPCR (6) |
| 8356 | CCTCGTCCTGCAGTTCATTC |  | 3’ primer |  |
| 8357 | ATGGCCGCTTTTCTGGATTC |  | 5’ primer | qPCR (7) |
| 8358 | CGATACCGTAAAGCACGAGG |  | 3’ primer |  |
| 8369 | TTTTGTCAAGACCGACCTGTCC |  | 5’ primer | qPCR (8) |
| 8370 | CTTCAGTGACAACGTCGAGC |  | 3’ primer |  |
| 8362 | CTTCCATGCCCGATGCACTT |  | 5’ primer | qPCR (CEN- HpaII) |
| 8361 | AAGTGCATCGGGCATGGAAG |  | 3’ primer |  |
| 8367 | ACCCAGTGTCTACAGGGTAATTTG |  | 5’ primer | qPCR (CEN- BstUI/) |
| 8368 | ACGAGGGGCATGAGGACCTCTG |  | 3’ primer |  |
| 8371 | TGTCCCCAGTCTCTTAGAG |  | 5’ primer | qPCR (CEN-Hpy99I) |
| 8372 | CTTGGTGAGTGAAGTAATG |  | 3’ primer |  |
